# Discovery of Novel Ligands for Cryptococcus neoformans

**DOI:** 10.64898/2026.03.05.709863

**Authors:** Teresa Rocha, Catarina Alves, Carla Lima, Lisa Sequeira, Fernanda Borges, Fernando Cagide, Sofia Benfeito

## Abstract

Fungal pathogens are an escalating global public health concern, particularly in the context of invasive and opportunistic infections. Cryptococcosis, primarily caused by Cryptococcus neoformans var. grubii, can manifest as acute, subacute, or chronic disease, affecting multiple organs and frequently leading to life-threatening meningitis in immunocompromised individuals. Given the limited antifungal therapeutic strategies and the emergence of resistance and toxicity-related constraints, the development of novel anti-cryptococcal agents remains an urgent priority.

In this study, a library of innovative hybrids (**5a-f**) based on the 3-hydroxypyridin-4(1H)-one scaffold was developed. Their antimicrobial activity was evaluated towards a panel of clinically relevant Gram-positive (methicillin-resistant *Staphylococcus aureus* – MRSA) and Gram-negative bacteria (*Escherichia coli*, *Klebsiella pneumoniae*, *Pseudomonas aeruginosa*, *Acinetobacter baumannii*), as well as fungal species *Candida albicans* and *Cryptococcus neoformans var. grubbi*. Cytotoxicity was assessed in HEK293 and HepG2 cell lines, and haemolytic profile was determined to evaluate safety. In addition, iron-chelating capacity and lipophilic properties were also investigated. All compounds formed stable complexes with iron(III) and were non-toxic at concentrations up to 25 μM. Lipophilicity studies showed that compounds in series 1 (**5a-c**) exhibited lower lipophilicity than those in Series 2 (**5d-f**), mainly due to the regioisomeric position of the hydroxyl group on the 2-methyl-4-pyridone scaffold; specifically, the C3-substitution pattern in Series 2 that enhances the hydrophobic character compared to the C5-substitution in Series 1. Fluorination further increased lipophilicity in both series. Notably, compounds **5c–5f** emerged as potent, selective, and non-toxic antifungal agents against *Cryptococcus neoformans* var. *grubii* (MIC < 16 µg/mL; CC50 > 32 µg/mL; HC10 > 32 µg/mL). Their distinct structural features appear to play a key role in antifungal selectivity, supporting the potential of these 3-hydroxypyridin-4(1H)-one-based hybrids as promising approach for the development of novel therapeutics for *cryptococcal meningitis*.

## 1. Introduction

Emerging fungal pathogens pose a growing threat to global public health. In response to this escalating burden, the World Health Organization has established a Fungal Priority Pathogens List, labelling *Cryptococcus neoformans* within the critical priority group [1, 2]. It is a ubiquitous fungal pathogen found worldwide, particularly in pigeon droppings and swampy environments. Infection is typically acquired through inhalation of airborne spores, which initially colonize the lungs. Pulmonary infection may lead to pneumonia, and in severe cases, the pathogen can disseminate via the bloodstream to the central nervous system (CNS), where it causes meningites (*cryptococcal meningitis*). *Cryptococcal meningitis* is a life-threatening fungal infection of the CNS caused by *Cryptococcus* species that invade the meninges and/or brain parenchyma, most commonly *Cryptococcus neoformans*. In 2020, fungal meningitis attributed to *C. neoformans* was associated with an estimated 112,000 deaths worldwide, underscoring its major global health impact [3, 4]. Complications due to *C. neoformans* infection and its treatment included acute renal impairment, elevated intracranial pressure requiring cerebrospinal fluid shunting, and vision loss. Clinically, patients typically present with fever, headache, neck stiffness, and visual disturbances. In immunocompetent individuals, infection is frequently mild or asymptomatic and is typically controlled by an effective immune response [5]. Conversely, severe disease occurs in the absence of adequate cell-mediated immunity, particularly in individuals with HIV/AIDS or in organ transplant recipients undergoing immunosuppressive therapy.

For the treatment of *Cryptococcal Meningitis*, only three antifungal agents are currently approved for clinical use: amphotericin B, flucytosine, and fluconazole (**Figure 1**) [6, 7]. Among these, amphotericin B (AmB), a polyene antifungal, remains the cornerstone of therapy for systemic infections caused by *Cryptococcus* infections due to its broad-spectrum activity and relatively low rates of resistance [8]. However, prolonged AmB therapy can cause serious adverse effects, including nephrotoxicity, hypokalemia, hypomagnesemia, and anemia [9]. The widespread and often prolonged use of antimicrobial agents drives the development of drug resistance. Likewise, the extensive and prolonged use of current antifungal therapies has contributed to the emergence of resistant fungal strains, thereby further undermining treatment efficacy. This growing threat underscores the urgent need for the development of novel and more effective antifungal agents.

**Figure 1.**
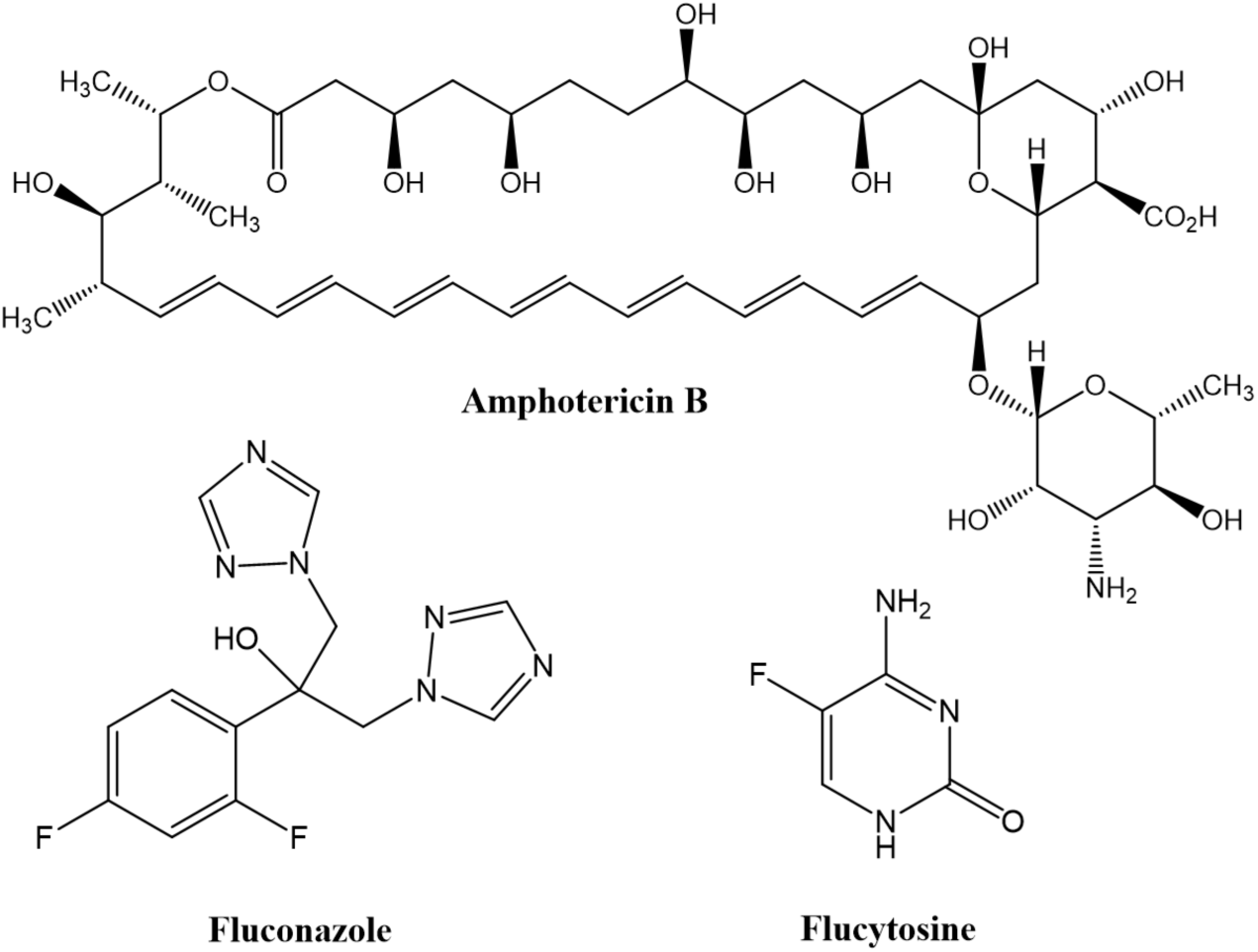
– Currently approved antifungal agents for the treatment of *Cryptococcal Meningitis*.

Although important advances have been made in antifungal therapy, the available therapeutic arsenal remains limited when compared to the broad range of antibiotics for bacterial infections. Expanding the antifungal pipeline is therefore both urgent and essential. Nevertheless, the discovery and development of new safe and effective antifungal drugs remain highly challenging, and fungal infections—despite their increasing global impact—continue to receive disproportionately limited attention and resources worldwide.

In recent years, fluorine-containing compounds, namely fluorinated cinnamic derivatives, have attracted growing attention as potential antimicrobial agents [10, 11]. Compared to their non-fluorinated counterparts, fluorinated derivatives often show lower minimum inhibitory concentration (MIC) values, along with improved selectivity and low cytotoxicity in preliminary evaluations [12–15]. The antimicrobial activity of these compounds is strongly influenced by both the number and the position of fluorine atoms on the aromatic ring. For instance, *para*-fluorination has been reported in some studies to enhance biological activity, whereas fluorination at other positions at other positions may reduce efficacy, depending on the specific biological target [16, 17]. These structure–activity differences are thought to arise from fluorine-induced modifications of physicochemical and electronic properties, which can influence key biological interactions. Proposed mechanisms include alterations in membrane permeability, enzyme inhibition mediated by electronic effects, and potential modulation of efflux pump activity.

In parallel with membrane and enzyme-targeting effects, disruption of microbial iron homeostasis represents another promising antimicrobial strategy [18]. Bacteria and fungi, like all living organisms, require iron as an enzyme cofactor to catalyze vital biological reactions. The uptake of iron by these unicellular organisms is accomplished through several mechanisms, including the production of small organic iron chelating molecules, called siderophores. These metabolites scavenge iron from the surrounding environment via specific receptors and enable its transport into the cytoplasm [19–21]. As iron uptake is crucial for microbial growth and virulence, modulation of iron homeostasis has emerged as a promising approach for antifungal therapy. Based on these assumptions, we developed a small library of compounds (**5a–f**) derived from 3-hydroxypyridin-4(1H)-one, a molecule known for its iron-chelating properties. Methyl groups were introduced at different positions on the heterocyclic core, which was subsequently conjugated to fluorinated cinnamic acid moieties through an alkyl linker. This preliminary and innovative study aims to develop hybrid antifungal agents with multiple mechanisms of action (**Figure 2**).

**Figure 2.**
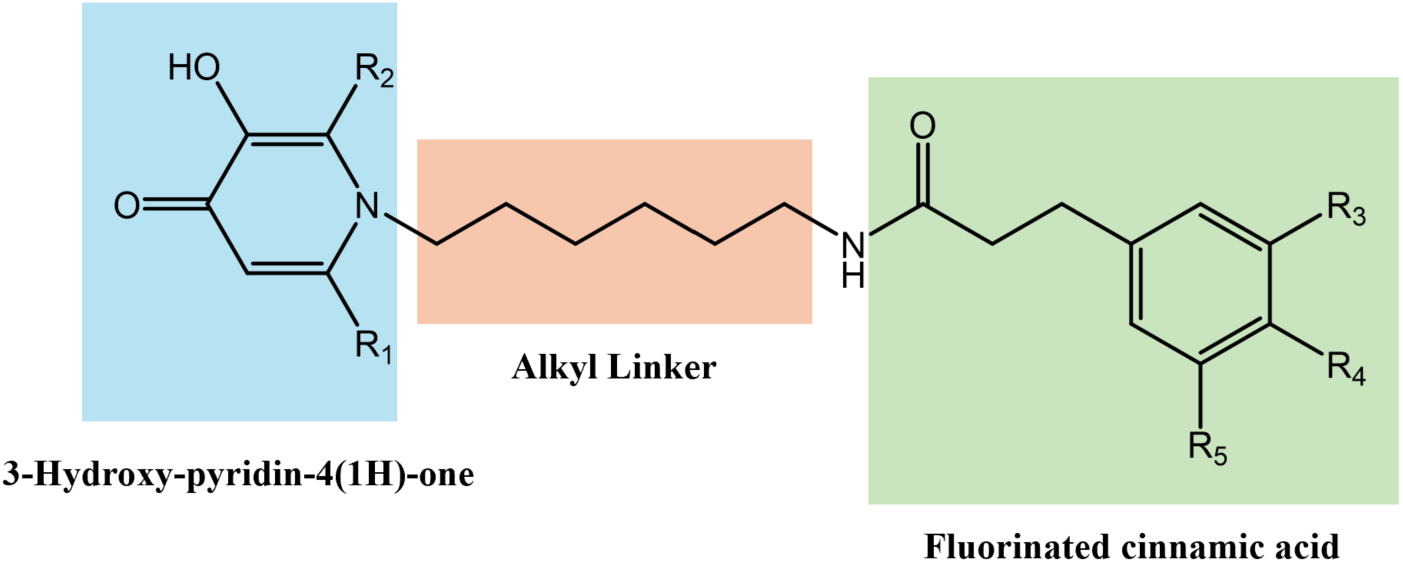
– Rational design of hybrids based on 3-hydroxypyridin-4(1H)-one.

## 2. Results and discussion

### 2.1 Chemistry

The 3-hydroxypyridin-4(1H)-one–based hybrids were synthesized through a five-step reaction sequence, as outlined in **Scheme 1**. Structurally, these compounds comprise a 3-hydroxypyridin-4(1H)-one core linked via a six-carbon alkyl spacer to a cinnamic acid moiety bearing one, two, or three fluorine atoms. The library was divided into two series according to the substitution pattern on the heterocyclic core. In Series 1, the methyl group is positioned adjacent to the hydroxyl group at the C5-position, whereas in Series 2, the methyl group is arranged relative to the hydroxyl group at the C3-position. This regioisomeric variation enables the evaluation of how the relative positioning of the methyl and hydroxyl substituents influences the physicochemical and biological properties of the hybrids. Within each series, structural diversity is achieved by varying the number of fluorine atoms on the cinnamic acid ring.

Briefly, in a one-pot, two-step process, the hydroxyl group of compounds **1a–b** was first protected with benzyl group using benzyl chloride (BnCl) in a basic medium, followed by the introduction of a protected amine with a six-carbon spacer, replacing the oxygen of the heteroaromatic ring (**Scheme 1**, step *a*). The *Boc* group of intermediates **2a-b** was removed with trifluoroacetic acid (**Scheme 1**, step *b*). Subsequently, various fluorinated cinnamic acids were introduced via two different methods, using either ethyl chloroformate or *N,N′*-dicyclohexylcarbodiimide (DCC) as shown in **Scheme 1**, step *c* or *d*, yielding compounds **4a-f**. Finally, O-deprotection of **4a-f** with trifluoroacetic acid afforded the final compounds **5a-f** (**Scheme 1**, step *e*).

**Scheme 1.**
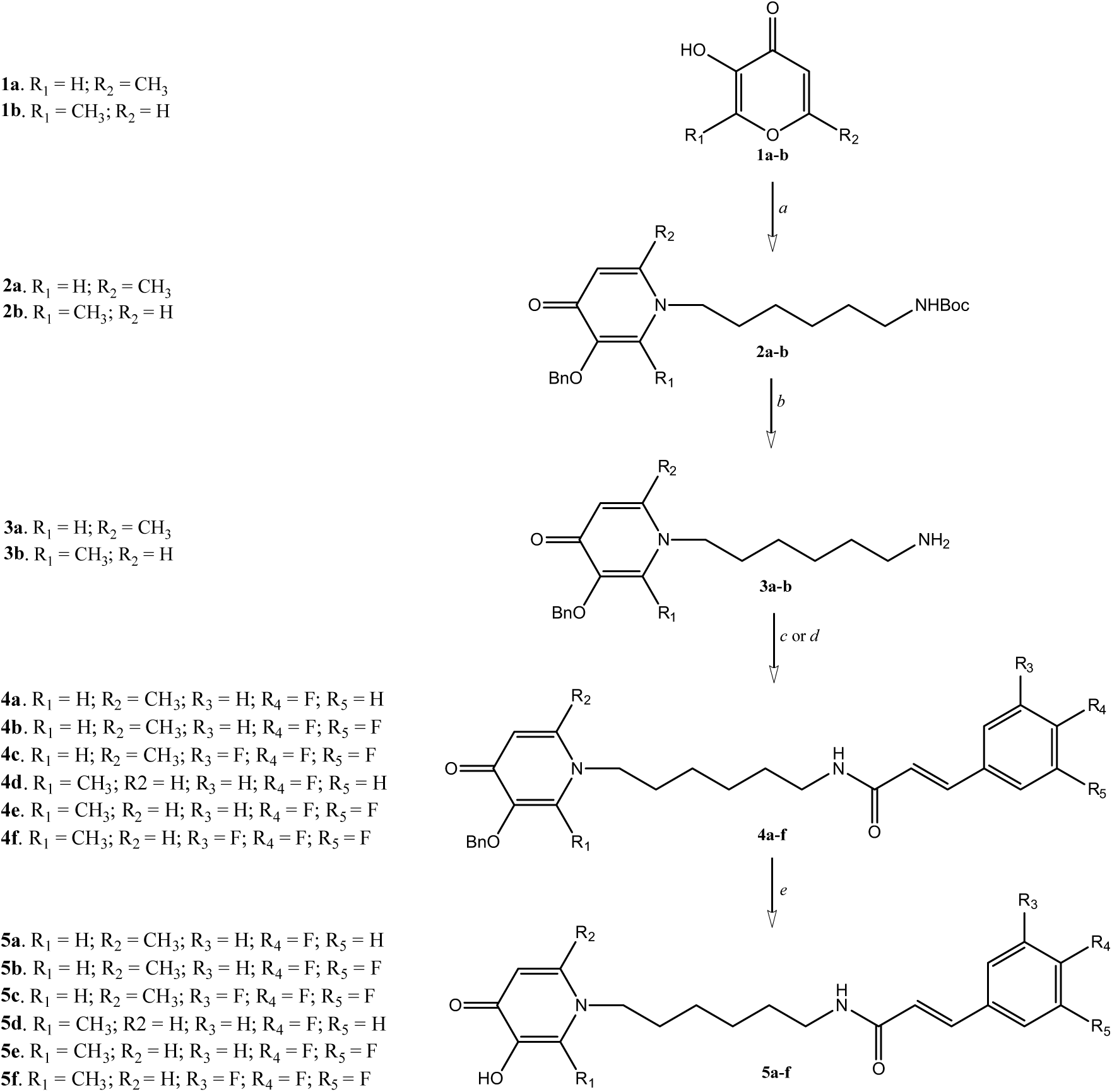
Synthetic strategy followed to obtain the hybrid compounds **5a-f**. Reagents and conditions: (a) BnCl, *tert*-butyl (6-aminohexyl)carbamate 80 °C, 28 h; (b) TFA, rt, 2 h; (c) Appropriate fluorinated cinnamic acids ((*E*)-3-(4-fluorophenyl)acrylic acid, (*E*)-3-(3,4-difluorophenyl)acrylic acid and (*E*)-3-(3,4,5-trifluorophenyl)acrylic acid), ClCOOEt, Et_3_N, DCM, rt, 24 h; (d) Appropriate fluorinated cinnamic acids ((*E*)-3-(4-;)acrylic acid, (*E*)-3-(3,4-difluorophenyl)acrylic acid and (*E*)-3-(3,4,5-trifluorophenyl)acrylic acid), DCC, NBSOH, Et_3_N, DCM, rt, 24 h; (e) TFA:Toluene (1:1), 65 °C, 24 h.

### 2.2 Antimicrobial screening

The *in vitro* antibacterial activity of the final compounds **5a–f** was assessed by CO-ADD (Community for Open Antimicrobial Drug Discovery). A preliminary high-throughput screening (HTS) was performed, with all compounds tested at a single concentration of 32 µg/mL against a variety of microbial strains, including 5 bacteria: *E. coli*, *K. pneumoniae* (MDR), *A. Baumannii*, *P. aeruginosa* and *S. aureus* (MRSA) and 2 fungi: *C. albicans* and *C. neoformans*. Vancomycin and colistin were used as positive references for Gram-positive and Gram-negative bacteria, respectively. For *C.albicans* and *C.neoformans*, fluconazole was used as a reference (**Table 1**). The HTS results were expressed based on growth inhibition (%) and Z-Score at a concentration of 32 µg/mL. Samples with inhibition values above 80% and Z-Score above |2.5| for either replicate (n =2 on different plates) were classified as active and selected for further evaluation. All hybrid compounds were found to be active. This criterion was crucial in identifying compounds requiring further dose-response confirmation. Then, the minimum inhibitory concentrations (MICs) of the active compounds were determined using the broth microdilution method against planktonic cells. An eight-point dose–response assay (32–0.25 µg/mL) was performed for each compound. The MIC (µg/mL) was defined as the lowest concentration at which growth was fully inhibited, corresponding to an inhibition of ≥ 80%. Colistin sulfate and Vancomycin hydrochloride were used as references for Gram-negative and Gram-positive bacteria, respectively, while Fluconazole served as the antifungal control against *C. albicans*.

**Table 1.** MIC values (µg/mL) of the hybrid compoundss **5a-f** as well as antibacterial and antifungal references against the selected microorganisms.

|  |  | MIC |  |  |  |  |  |  |
| --- | --- | --- | --- | --- | --- | --- | --- | --- |
| Series | Compounds | *S. aureus* | *E. coli* | *K. pneumoniae* | *P. aeruginosa* | *A. baumannii* | *C. albicans* | *C. Neoformans* |
| 1 | **5a** | >32 | >32 | >32 | >32 | >32 | >32 | >32 |
|  | **5b** | >32 | >32 | >32 | >32 | >32 | >32 | >32 |
|  | **5c** | 32 | >32 | >32 | >32 | >32 | >32 | 16 |
| 2 | **5d** | 32 | >32 | >32 | >32 | >32 | >32 | 16 |
|  | **5e** | >32 | >32 | >32 | >32 | >32 | >32 | 4 |
|  | **5f** | >32 | >32 | >32 | >32 | >32 | >32 | 4 |
|  | **Colistin** | >32 | 0.125 | 0.25 | 0.25 | 0.25 | n.d | n.d |
|  | **Vancomycin** | 1 | >32 | >32 | >32 | >32 | n.d | n.d |
|  | **Fluconazole** | n.d | n.d | n.d | n.d | n.d | 0.125 | 8 |

Standard strains: *S. aureus* ATCC 43300; *E. coli* ATCC 25922, *K. pneumoniae* ATCC 70060, *P. aeruginosa* ATCC 27853, *A. baumannii* ATCC 19606, *C. albicans* ATCC 90028, and *C. neoformans* var. *grubii* H99 ATCC 208821. n.d.: not determined

To streamline the analysis, the compounds were organized into two series. In series 1, the methyl group is positioned adjacent to the hydroxyl group at the C5 of the heterocyclic core, whereas in series 2, it is located relative to the hydroxyl group at the C3-position, enabling evaluation of the impact of regioisomeric substitution on antimicrobial activity. In the antimicrobial dose–response assays, all hybrid compounds were confirmed as hits (**Table 1**), validating the initial high-throughput screening results. Notably, compounds **5c–5f** showed pronounced antifungal activity against the *C. neoformans* strain, with MIC values ≤ 16 µg/mL. Among them, compounds **5e** and **5f** emerged as the most promising candidates, exhibiting superior potency compared to the reference antifungal Fluconazole. These findings highlight the strong therapeutic potential of this hybrid scaffold and underscore its relevance for the development of new anti-cryptococcal agents.

### 2.3 Cytotoxicity and haemolytic screening

The toxicity profile of compounds **5a–f** was investigated by assessing their effect on the viability of the mammalian HEK293 cell line, derived from human embryonic kidneys, and their potential to induce erythrocyte haemolysis. Cytotoxicity was expressed as the CC_50_ (concentration causing 50% reduction on cell viability) and HC_10_ (concentration inducing 10% haemolysis of human red blood cells). The corresponding values are summarized in **Table 2**. Tamoxifen and Melittin were used as positive controls for cytotoxicity and haemolytic activity, respectively. Importantly, none of the tested compounds exhibited detectable cytotoxicity in HEK293 cells, and no haemolytic activity against human red blood cells was detected at concentrations up to 32 µg/mL, suggesting a favorable preliminary safety profile.

**Table 2.** Cytotoxicity in HEK293 cells (CC_50_ in µg/mL) and haemolytic activity in RBC (HC_10_ in µg/mL) of the compounds.

| Series | Compounds | HEK293 (CC<sub>50</sub>)<sup>a</sup> | RBC (HC<sub>10</sub>)<sup>b</sup> |
| --- | --- | --- | --- |
| 1 | 5a | >32 | >32 |
|  | 5b | >32 | >32 |
|  | 5c | >32 | >32 |
| 2 | 5d | >32 | >32 |
|  | 5e | >32 | >32 |
|  | 5f | >32 | >32 |
|  | Tamoxifen | 9 | - |
|  | Mellitin | - | 2.7 |

<sup>a</sup>CC<sub>50</sub>: concentration at 50 % cytotoxicity in human embryonic kidney (HEK-293) cells.
<sup>b</sup>HC<sub>10</sub>: concentration at 10% haemolytic activity in human red blood (RBC) cells.

The cytotoxicity of compounds **5a–f** was further evaluated in HepG2 cells, a human hepatocellular carcinoma cell line commonly used for early hepatotoxicity screening. This model is widely employed in early-stage drug development to identify and eliminate potentially hepatotoxic compounds prior to *in vivo* studies. The cell viability was assessed 24 hours after exposure with six different concentrations (0.1, 0.5, 1, 5, 10 and 25 µM) using the resazurin reduction (**Figure 2A**) and neutral red (NR) uptake assays (**Figure 2B**), to select the non-cytotoxic concentration ranges of the compounds.

In the resazurin assay, none of the compounds (**5a–f**) induced cytotoxicity at any tested concentration after 24 h exposure (**Figure 2A**). In the NR uptake assay (**Figure 2B**), compound **5a** exhibited no cytotoxic effects across the tested concentration range. The compounds **5b and 5c** showed modest reductions in NR uptake at concentrations above 10 µM, with viability decreasing to 87% and 86% for **5b**, and to 91% and 84% for **5c**, at 10 and 25 µM, respectively. Within series 2, compound **5d**, bearing a single fluorine atom, showed no evidence of cytotoxicity up to 25 µM. In contrast, compound **5e**, induced a moderate but statistically significant reduction in NR uptake at concentrations ≥10 µM (90% viability at 10 µM and 89% at 25 µM), while compound **5f**, containing three fluorine atoms, caused only a slight decrease at 25 µM (89% viability).

**Figure 2.**
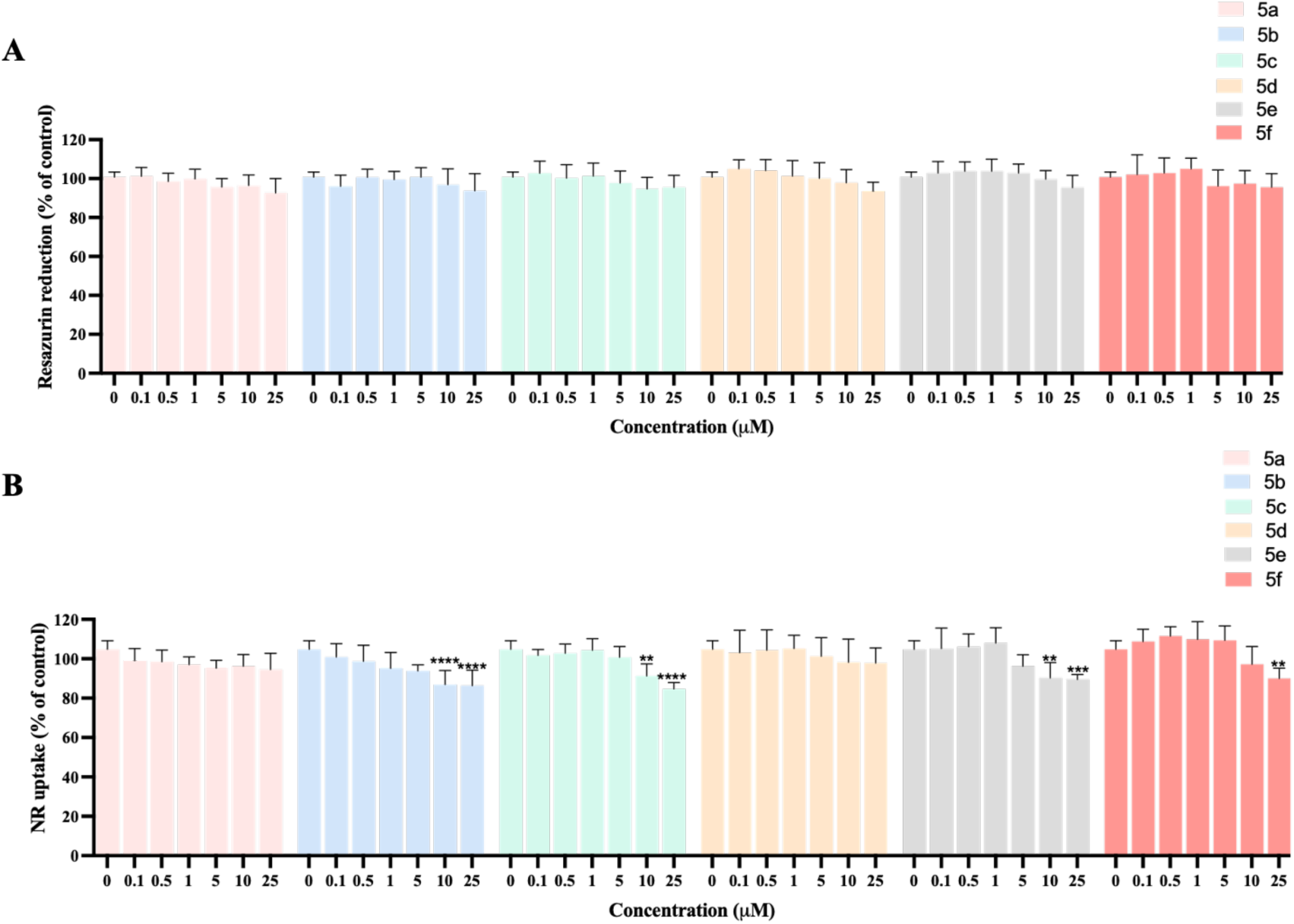
Cytotoxicity of compounds **5a-f** in HepG2 cells was assessed by the resazurin. (A) and neutral red (B) methods after 24 hours of treatment. The results are expressed as the mean and standard deviation (mean ± SD) of three independent tests carried out in triplicate. Statistical comparisons were carried out using the parametric one-way ANOVA method, following the Dunnett’s multiple comparison test [** p ˂ 0.01, *** p ˂ 0.001, **** p ˂ 0.0001, vs. CTRL (0 μM)].

Importantly, despite the slight decreases in NR uptake observed for some derivatives, all compounds showed high cell viability, and the consistent results across both resazurin reduction and neutral red uptake assays—assessing metabolic activity and lysosomal function, respectively—confirm a robust safety profile for compounds **5a–f** in HepG2 cells at concentrations up to 25 µM after 24 hours of exposure.

### 2.4 Determination of lipophilicity

The determination of lipophilicity is a key parameter in drug discovery, as it provides insight into the overall drug-likeness of a compound. Lipophilicity strongly influences the passive diffusion across cell membranes and biological barriers, thereby impacting pharmacological potency, bioavailability, and toxicity. The lipophilicity of compounds **5a-f** was experimentally determined using Chromatographic Hydrophobicity Index (CHI) at pH 2.6, using a fast-gradient HPLC approach. The CHI values were calculated by converting the chromatographic retention times (t_r_) of the compounds into hydrophobicity indices using calibration curves generated from a standard mixture of reference compounds with known hydrophobicity. The reference compounds utilized was theophylline, paracetamol, caffeine, benzimidazole, colchicine, carbamazepine, indole, propiophenone, butyrophenone and valerophenone (**Figure 5**). The results obtained for compounds **5a–f** are summarized in **Table 3**.

**Table 3.**
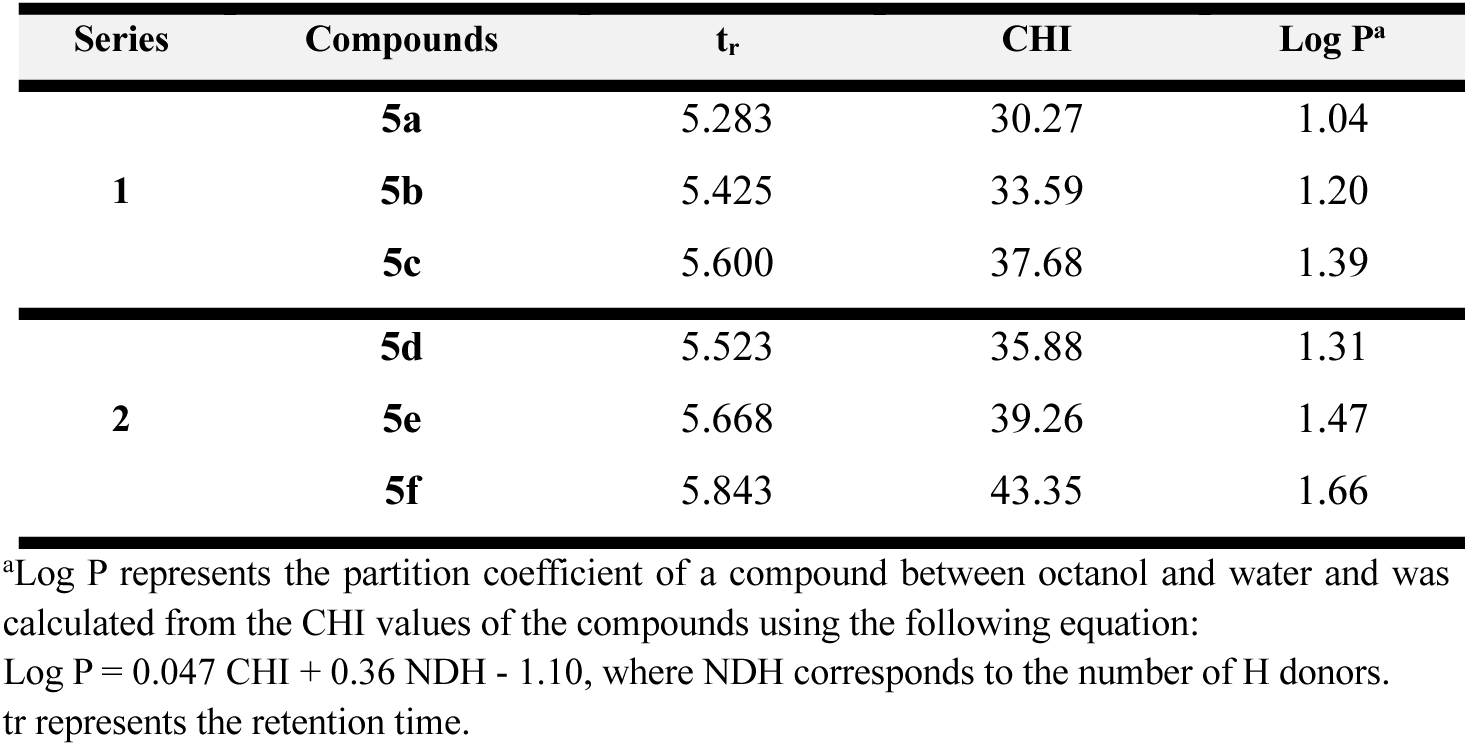
Experimental CHI values at pH 2.6 of compounds **5a-f**.

| Series | Compounds | $t_r$ | CHI | Log P<sup>a</sup> |
| --- | --- | --- | --- | --- |
| 1 | **5a** | 5.283 | 30.27 | 1.04 |
|  | **5b** | 5.425 | 33.59 | 1.20 |
|  | **5c** | 5.600 | 37.68 | 1.39 |
| 2 | **5d** | 5.523 | 35.88 | 1.31 |
|  | **5e** | 5.668 | 39.26 | 1.47 |
|  | **5f** | 5.843 | 43.35 | 1.66 |

<sup>a</sup>Log P represents the partition coefficient of a compound between octanol and water and was calculated from the CHI values of the compounds using the following equation: $\text{Log P} = 0.047 \text{ CHI} + 0.36 \text{ NDH} - 1.10$ , where NDH corresponds to the number of H donors. $t_r$ represents the retention time.

A comparative analysis of the two series revealed a clear influence of substitution pattern of hybrid compounds on lipophilicity. Compounds from series 2 (**5d-f**) showed higher CHI values than those in series 1 (**5a-c**). This increase can be attributed to the presence of a methyl group positioned relative to the hydroxyl at C3 of the heterocyclic core, a structural arrangement that enhances hydrophobic character and results in higher Log P values (**5d**: 1.31; **5e**:1.47; **5f**: 1.66). In addition to the effect of methyl positioning, fluorination also contributed to increased lipophilicity in both series. An increase of CHI and corresponding Log P values was observed with the introduction of additional fluorine atoms on the cinnamic moiety. For example, comparison of **5c** and **5f** (Log P = 1.39 and 1.66, respectively) highlights the cumulative impact of hybrids substitution pattern.

Together, these findings demonstrate that both the position of the methyl group on the heterocyclic core and the degree of fluorination on the cinnamic fragment synergistically modulate lipophilicity. Moreover, the overall structural features of compounds **5a–f** suggest favorable physicochemical characteristics that may promote passive diffusion across cellular membranes.

### 2.5 Evaluation of iron (III) chelation

Iron acquisition is a key factor of microbial pathogenesis and represents an attractive target for antifungal therapy. In *C. neoformans*, iron availability directly regulates the elaboration of the polysaccharide capsule, a major virulence determinant of the fungus. To evaluate the Fe(III)-chelating capacity of compounds **5a–f**, their UV–Vis spectra were recorded both in the absence and presence of the metal ion. As shown in **Figure 3**, the spectra of the compounds (40 µM) in phosphate buffer were compared with those obtained after co-incubation with Fe(III) chloride (20 µM), providing insight into their ability to interact with iron.

**Figure 3.**
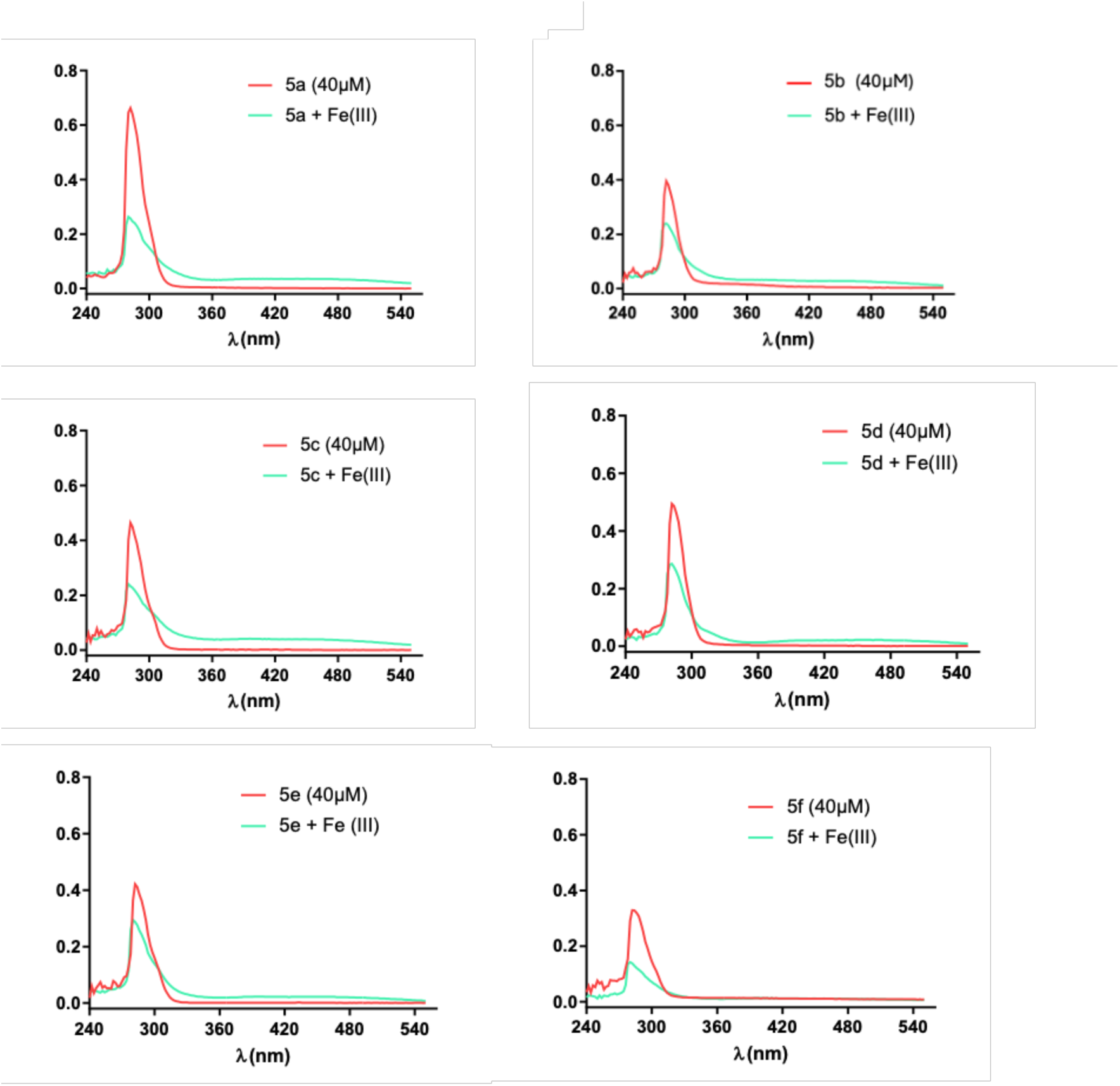
UV–Vis spectra of compounds **5a–f** (40 µM) recorded in the absence and presence of iron (III) chloride (20 µM).

Next, the stoichiometry of the Fe(III)–compound complex was determined using the molar ratio method. In these experiments, the concentration of each test compound was fixed, while the concentration of Fe(III) chloride was gradually increased from 0 to 100 µM (**Figure 4**). The results confirmed that all compounds are capable of forming complexes with Fe(III). The stoichiometry of the Fe(III)–compound complexes was determined by varying the concentration of iron(III) chloride.

**Figure 4.**
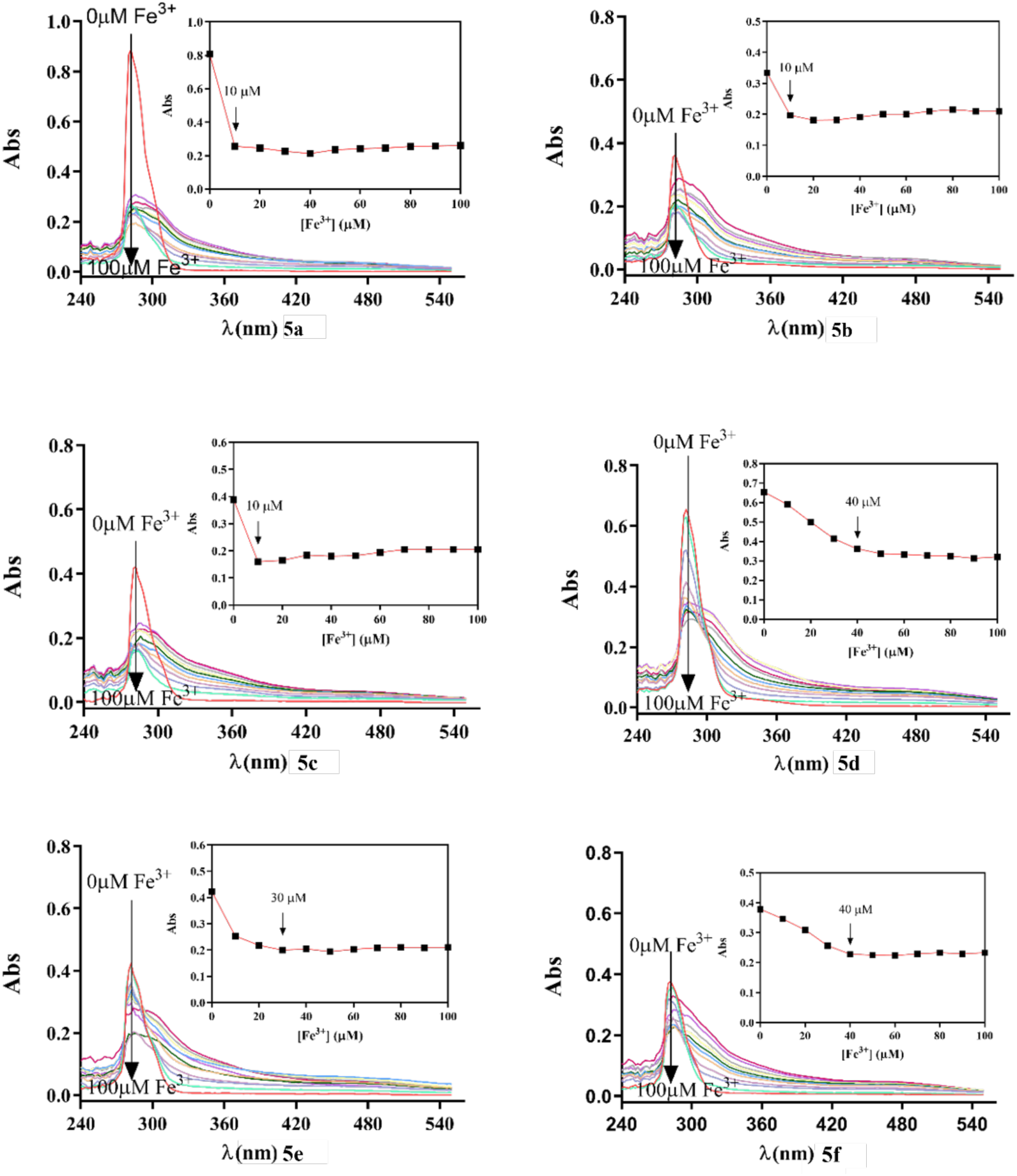
UV-Vis titration of compounds **5a-f** with iron (III) in PBS at room temperature.

Compounds from series 1, in which the methyl group is positioned adjacent to the hydroxyl group at C5 of the heterocyclic core, displayed a 0.25:1 stoichiometry. In contrast, compounds **5d** and **5f** exhibited a 1:1 Fe³⁺/ligand stoichiometry, whereas compound **5e** from the same series showed a 0.75:1 ratio. These findings indicate that compounds **5d** and **5f** have a greater capacity to form complexes with Fe(III), likely due to the presence of a methyl group positioned relative to the hydroxyl at C3 of the heterocyclic core, combined with one (compound **5d**) or three (compound **5f**) fluorine atoms on the cinnamic acid moiety — structural features that may enhance their ability to coordinate Fe(III).

### 2.6 Structure-activity-toxicity-lipophilicity-chelation relationships

By correlating the structure–activity and toxicity data (**Tables 1** and **2**; **Figure 2**), it can be concluded that derivatives from **series 2** and compound **5c** from **series 1** demonstrated a meaningful antifungal activity against *C. neoformans* var. *grubii* (**Table 1**). A clear correlation was observed between antifungal activity against *C. neoformans* var. *grubii*. and the structural features of the compounds, with both the position of the methyl group on the heterocyclic core and the extent of fluorination on the cinnamic moiety playing a significant role. Notably, compounds with a methyl group positioned relative to the hydroxyl at C3 of the heterocyclic core (**5d–5f),** exhibited the most remarkable antifungal activity, emphasizing the key role of this substitution pattern in antifungal activity. In addition, compounds containing two or three fluorine atoms exhibited lower MIC values compared to the mono-fluorinated derivative (**5e–5f** vs. **5d**, respectively), suggesting that increased fluorination enhances antifungal activity.

Overall, these results suggest that optimal antifungal activity is achieved through a combination of a methyl group positioned relative to the hydroxyl at C3 of the heterocyclic core and selective fluorine substitution on the cinnamic moiety.

Based on the cytotoxicity data obtained in HEK293 and HepG2 cells, none of the compounds exhibited significant cytotoxic effects (**Table 2**, **Figure 2**). Likewise, assessment of haemolytic activity against human red blood cells (RBCs) confirmed that all compounds were non-haemolytic (**Table 2**). This favorable safety profile, combined with the compounds’ predominantly lipophilic character—which is further enhanced by a methyl group positioned relative to the hydroxyl at C3 of the heterocyclic core and by an increasing number of fluorine atoms—suggests that these compounds are well-suited to cross cell membranes, supporting their potential as effective antifungal agents. Furthermore, UV–Vis and molar ratio studies confirmed that all compounds are capable of forming complexes with Fe(III). Notably, compounds **5d** and **5f** exhibited a greater capacity to chelate Fe(III), likely due to the combination of the methyl group positioned relative to the hydroxyl at C3 of the heterocyclic core and the presence of one (**5d**) or three (**5f**) fluorine atoms on the cinnamic acid moiety—structural features that appear to enhance iron coordination.

## 3. CONCLUSIONS

All novel 3-hydroxypyridin-4(1H)-one–based hybrids (**5a–f**) were successfully synthesized through a five-step synthetic sequence. These compounds were rationally designed as two distinct series differing in the substitution pattern on the heterocyclic core. In Series 1, the methyl group is positioned adjacent to the hydroxyl group, whereas in Series 2, it is arranged relative to the hydroxyl at the C3-position. This structural variation allowed a systematic assessment of how the relative positioning of the methyl group modulates the physicochemical and biological properties of the hybrid compounds. Additionally, within each series, modulation of the fluorination pattern on the cinnamic acid fragment provided further structural diversity, enabling the exploration of structure–activity–property relationships.

Compounds **5c–5f** demonstrated pronounced antifungal activity against *C. neoformans* (MIC ≤ 16 µg/mL), with **5e** and **5f** showing superior potency compared to fluconazole, highlighting their therapeutic potential. Importantly, all compounds exhibited a favorable safety profile, with no detectable cytotoxicity in HEK293 and HepG2 cells, and no haemolytic activity up to 32 µg/mL. Furthermore, the analysis of both series indicates that lipophilicity is modulated by the position of the methyl group relative to the hydroxyl at C3 of the heterocyclic core and the level of fluorination on the cinnamic fragment. Series 2 compounds (**5d–f**) showed higher CHI and Log P values than Series 1 (**5a–c**), reflecting the hydrophobic contribution of the methyl group relative to the hydroxyl at C3. This effect was further enhanced by additional fluorine atoms on the cinnamic moiety, demonstrating a synergistic influence of core substitution and fluorination on the overall lipophilic character. Finally, the Fe(III) complexation studies revealed that compounds **5d** and **5f** exhibit the greatest capacity to coordinate iron, forming 1:1 Fe³⁺/ligand complexes, whereas **Series 1** compounds and **5e** show lower stoichiometries. This enhanced chelating ability appears to result from the combined effect of the methyl group positioned relative to the hydroxyl at C3 of the heterocyclic core and the presence of fluorine atoms on the cinnamic acid moiety.

Overall, these findings underscore the importance of 3-hydroxypyridin-4(1H)-one hybrids as a promising approach for the development of novel anti-cryptococcal agents.

## 4. Experimental section

### 4.1 Chemistry

#### 4.1.1 Reagents and apparatus

Starting materials, reagents, and solvents were obtained from commercial suppliers and were used without any further purification. Deionized water was used for all syntheses. All reactions were followed by thin-layer chromatography (TLC) using aluminum plates coated with silica gel 60 F254 (Merck). For analytical control, the mobile phase used was dichloromethane/methanol in several proportions. The chromatograms obtained were revealed using ultraviolet (UV) light (Vilber corporation) with a wavelength of 254 and/or 365 nm and/or an aqueous solution of 10 % (w/v) iron chloride and/or a 0.2 % (w/v) ninhydrin solution. The crude products were purified by flash column chromatography on silica gel 60 Å (0.040–0.063 mm, Carlo Erba Reactifs) and/or by recrystallization. The specific elution systems employed for flash chromatography are described individually for each compound. Finally, anhydrous sodium sulfate (Na_2_SO_4_) (Honeywell FlukaTM) was used to dry the organic phases.

NMR data were acquired on a Bruker Avance III 400 NMR spectrometer, at room temperature, operating at 400.15 MHz for ^1^H and 100.62 MHz for ^13^C and DEPT135 (Distortionless Enhancement by Polarization Transfer). Tetramethylsilane (TMS) was used as internal reference; chemical shifts (δ) were expressed in ppm and coupling constants (J) were given in Hz. DEPT135 values were included in ^13^C NMR data (underline values). The absorbance readings were used in different microplate readers, the Biotek Synergy HT and EpochTM Microplate Spectrophotometer (BioTek), and the BRANDplates® pureGradeTM and the Greiner bio-one UVSTAR® plates were used.

#### 4.1.2 Synthesis

##### 4.1.2.1. General procedures for the synthesis of intermediate **2a-b**

Compounds **2a-b** were obtained following procedure **A**.

**Procedure A**. The appropriate compounds **1a-b** (1 mmol) and sodium hydroxide (1.1 mmol) were dissolved in methanol (1.2 ml) under stirring at 60 °C. Then benzyl chloride (1.1 mmol) was added and the temperature was raised to 80 °C. After 4 hours, distilled water (0.6 mL) and tert-butyl(6-aminohexyl)carbamate (3.6 mmol) were added, and the pH was maintained at 8-9. After 24 hours, the solvent was evaporated, and an extraction was carried out. The organic phase was dried with anhydrous Na_2_SO_4_, Filtered, and evaporated. Finally, the compound was purified by silica gel flash column chromatography with DCM/MeOH 9.5:0.5 as eluent. The experimental procedure was adapted from the literature [22].

***Tert-butyl (6-(5-(benzyloxy)-2-methyl-4-oxopyridin-1 (4H)-yl) hexyl) carbamate (2a)***. Procedure A. η = 69 %. ^1^H RMN (400 MHz, CDCl_3_): δ = 7.43-7.37 (m; 2H; CH (2’/6’)); 7.37-7.27 (m; 3H; CH (3’/4’/ 5’)); 6.87 (s; 1H; CH (2)); 6.34 (s; 1H; CH (5)); 5.18 (s; 2H; CH_2_ (7)); 3.65 (t; *J* = 7.5 Hz; 2H; CH_2_ (a)); 3.09 (q; *J* = 6.8; 6.0 Hz; 2H; CH_2_ (f)); 2.24 (s; 3H; CH_3_ (6)); 1.56 (p; *J* = 7.5 Hz; 2H; CH_2_ (b)); 1.45-1.37 (m; 11H; (CH_3_)_3_; (OC=O) and CH_2_ (e)); 1.30-1.21 (m; 4H; CH_2_ (c/d)).^13^C RMN (101 MHz, CDCl_3_): δ = 173.1 (C (4)); 156.0 (C (g)); 147.4 (C (3)); 144.8 (C (6)); 136.9 (C (1’)); <u>128.5</u> (C (3’/5’)); <u>128.1</u> (C (4’)); <u>128.0</u> (C (2’/6’)); <u>118.5</u> (C (2)); <u>77.2</u> ((5)); <u>71.9</u> ((7)); <u>53.1</u> (C (a)); 40.3 (C (h)); <u>30.5</u> (C (f)); <u>29.9</u> (C (b)); <u>28.4</u> (C (i/j/k)); <u>26.2</u> (C (c)); <u>25.9</u> (C (d/e)); <u>18.9</u> (C (8)).

***Tert-butyl (6-(3-(benzyloxy)-2-methyl-4-oxopyridin-1 (4H)-yl) hexyl) carbamate (2b).*** Procedure A. η = 53 %. ^1^H RMN (400 MHz, CDCl_3_): δ = 7.43-7.37 (m; 2H; CH (2’/6’)); 7.34-7.28 (m; 3H; CH (3’/4’/5’)); 7.16 (d; *J* = 7.5 Hz; 1H; CH (6)); 6.40 (d; *J* = 7.5 Hz; 1H; CH (5)); 5.21 (s; 2H; CH_2_ (7)); 4.56 (bs; 1H; NH); 3.71 (t; *J* = 7.6 Hz; 2H; CH_2_ (a)); 3.10 (q; *J* = 6.7 Hz; 2H; CH_2_ (f)); 2.07 (s; 3H; CH_3_ (2)); 1.60 (p; *J* = 6.9; 6.2 Hz; 2H; CH_2_ (b)); 1.43 (s; 9H; CH_3_ (OC=O)); 1.37-1.23 (m; 6H; CH_2_ (c/d/e).^13^C RMN (101 MHz, CDCl_3_): δ = 173.3 (C (4)); 156.0 (C (<u>g</u>)); 146.1 (C (3)); 140.6 (C (2)); <u>138.1</u> (C (6)); 137.6 (C (1’)); <u>129.2</u> (C (3’/5’)); <u>128.2</u> (C (2’/6’)); <u>127.9</u> (C (4’)); <u>117.3</u> (C (5)); <u>79.2</u> (C (h)); <u>72.9</u> (C (7)); <u>53.7</u> (C (a)); <u>40.2</u> (C (f)); <u>30.7</u> (C (b)); <u>29.9</u> (C (c)); <u>28.4</u> (C (i/ j/k)); <u>26.2</u> (C (e)); <u>26.0</u> (C (d); <u>12.4</u> (C (8)).

##### 4.1.2.2. General procedures for the synthesis of intermediate **3a-b**

Compounds **3a-b** were obtained following procedure **B**.

**Procedure B**. The compound **2a-b** (1 mmol) was dissolved in dichloromethane (3.3 mL) under stirring in an ice bath. Trifluoroacetic acid (TFA) (0.8 mL) was then added. After 2 hours, TFA was removed by vacuum and an extraction was carried out with dichloromethane distilled water and a 5% NaOH solution. The organic phase was dried with anhydrous Na_2_SO_4_ filtered, and concentrated. The experimental procedure was adapted from the literature [23].

***1-(5-aminopentyl)-5-(benzyloxy)-2-methylpyridin-4(1H)-one (3a).*** Procedure B. η = 61 %. ^1^H RMN (400 MHz, CDCl_3_): δ = 7.43-7.38 (m; 2H; CH (2’/6’)); 7.36-7.27 (m; 3H; CH (3’/4’/5’)); 6.86 (s; 1H; CH (2)); 6.32 (s; 1H; CH (5)); 5.18 (s; 2H; CH (7)); 3.65 (t; *J* = 7.5 Hz; 2H; CH_2_ (a)); 2.67 (t; *J* = 6.9 Hz; 2H; CH_2_ (f)); 2.24 (s; 3H; CH_3_ (6)); 1.64-1.43 (m; 6H; CH (b/c/e)); 1.49-1.36 (m; 2H; CH (d)); 1.35-1.21 (m; 2H; NH_2_). ^13^C RMN (101 MHz, CDCl_3_): δ = 173.4 (C (4)); 147.4 (C (3)); 144.7 (C (6)); 137.0 (C (1’)); <u>128.5</u> (C (3’/5’)); <u>128.2</u> (C (4’)); <u>128.0</u> (C (2’/6’)); <u>127.9</u> (C (2)); <u>118.6</u> (C (5)); <u>71.9</u> (C (7)); <u>53.1</u> (C (a)); <u>41.9</u> (C (f)); <u>33.3</u> (C (e)); <u>30.5</u> (C (b)); <u>26.5</u> (C (c)); <u>26.2</u> (C (d)); <u>18.9</u> (8)).

***1-(6-aminohezyl)-3-(benzyloxy)-2-methylpyridin-4(1H)-one (3b).*** Procedure B. η = 45%. ^1^H RMN (400 MHz, CDCl_3_): δ = 7.44-7.36 (m; 2H; CH (2’/6’)); 7.35-7.28 (m; 3H; CH (3’/4’/5’)); 7.18 (d; *J* = 7.5 Hz; 1H; CH (6)); 6.41 (d; *J* = 7.5 Hz; 1H; CH (5)); 5.21 (s; 2H; CH_2_ (7)); 3.73 (t; *J* = 7.6 Hz; 2H; CH_2_ (a)); 2.68 (t; *J* = 6.9 Hz; 2H; CH_2_ (f)); 2.08 (s; 3H; CH_3_ (2)); 1.68-1.56 (m; 2H; CH_2_ (b)); 1.48-1.38 (m; 2H; NH_2_); 1.38-1.21 (m; 6H; CH_2_ (c/d/e)). ^13^C RMN (101 MHz, CDCl_3_): δ = 173.3 (C (4)); 146.1 (C (3)); 140.7 (C (2)); <u>138.2</u> (C (6)); 137.6 (C (1’)); <u>129.1</u> (C (3’/5’)); <u>128.2</u> (C (2’/6’)); <u>127.9</u> (C (4’)); <u>117.3</u> (C (5)); <u>72.9</u> (C (7)); <u>53.8</u> (C (a)); <u>41.9</u> (C (f)); <u>32.2</u> (C (e)); <u>30.7</u> (C (b)); <u>26.4</u> (C); <u>26.2</u> (C (d)); 12.4 (C (8)).

##### 4.1.2.3. General procedures for the synthesis of intermediate **4a-f**

The fluorinated cinnamic amides **4a** and **4d** were obtained according to procedure **C2,** and the fluorinated cinnamic amides **4b**, **4c**, **4e,** and **4f** were obtained according to procedure **C1**.

**Procedure C1**. The appropriate fluorinated cinnamic acids (1 mmol) were dissolved in dichloromethane (7.8 mL), and then triethylamine (2 mmol) was added in an ice bath and stirred for 5 min. Ethyl chloroformate (2 mmol) was added. After 2 hours, once the intermediate formed compounds **3a-b** (2 mmol) were added to the reaction mixture in an ice bath. After 24 hours, the mixture was purified by column chromatography using a DCM/MeOH 9.5:0.5 as eluent. The experimental procedure was adapted from the literature [24].

**Procedure C2**. The appropriate fluorinated cinnamic acids (1 mmol), DCC (1.2 mmol), and N-Hydroxysuccinimide (NHS) (0.2 mmol) were dissolved in dichloromethane (12 mL). After 5 min. Et_3_N (4 mmol) was added in an ice bath and stirred. After 2 hours, the reaction was controlled, and intermediates **3a-b** (1 mmol) dissolved in DCM were added. The reaction was left to stir overnight. After reaction completion, the compound obtained was purified by column chromatography using DCM/MeOH 9.5:0.5 as eluent. The experimental procedure was adapted from the literature [25].

***(E)-N-(6-(5-(benzyloxy)-2-methyl-4-oxopyridin-1(4H)-yl)hexyl)-3-(4-fluorophenyl)acrylamide (4a)***. Procedure C2. η = 26 %.^1^H RMN (400 MHz, CDCl_3_): δ = 7.60 (t. *J* = 5.7. 1H; NH_2_); 7.53 (d; *J* = 15.7 Hz; 1H; CH (β)); 7.46-7.37 (m; 2H; CH (2’’/6’’)); 7.37-7.33 (m; 2H; CH (2’/6’)); 7.37-7.33 (m; 3H; CH (3’ – 5’)); 6.98-6.93 (m; 2H; CH (3’’/5’’)); 6.56 (d; *J* = 15.7 Hz; CH (α)); 6.26 (s; 1H; CH (2)); 5.29 (s; 1H; CH (5)); 5.08 (s; 2H; CH (7)); 3.68-3.59 (m; 2H; CH (a)); 3.35-3.26 (m; 2H; CH (f)); 2.18 (s; 3H; CH_3_ (6)); 1.60-1.43 (m; 4H; CH (b/e)); 1.37-1.25 (m; 2H; CH (c)); 1.24-1.16 (m; 2H; CH (d)). ^13^C RMN (101 MHz, CDCl_3_): δ = 172.9 (C (4)); 166.3 (C(g)); 163.2 (d; *J*_CF_ = 249.6 Hz C (4’’); 147.5 (C (3)); 145.2 (C (6)); <u>138.5</u> (C (β)) 136.5 (C (1’)); 131.5 (d; *J* = 3.4 Hz; C (1’’)); <u>129.4</u> (d; *J* = 8.2 Hz; C (2’’/6’’)); <u>128.5</u> (C (3’/5’)); <u>128.1</u> (C (4’)); <u>127.9</u> (C (2’/6’)); <u>127.6</u> (C (2)); <u>121.6</u> (C (α)); <u>117.9</u> (C (5)); <u>115.7</u> (d; *J* = 21.8 Hz; C (3’’/5’’)); <u>71.7</u> (C (7)); <u>53.2</u> (C (a)); <u>39.3</u> (C (f)); <u>30.3</u> (C (e)); <u>29.2</u> (C (b)); <u>26.3</u> (C (c)); <u>25.8</u> (C (d)); <u>18.8</u> (C (8)).

***(E)-N-(6-(5-(benzyloxy)-2-methyl-4-oxopyridin-1(4H)-l)hexyl)-3-(3.4 difluorophenyl)acrylamide (4b).*** Procedure C1. η = 25 %. ^1^H RMN (400 MHz, CDCl_3_): δ = 7.54-7.39 (m; 5H; CH (3’-5’/6’’/β)); 7.36-7.21 (m; 4H; CH (2’/6’/2’’/5’’)); 6.56 (d; *J* = 15.7 Hz; CH (α)); 6.35 (s; CH (2)); 5.48 (s; CH (5)); 5.07 (s; 2H; CH_2_ (7)); 3.96-3.87 (m; 2H; CH_2_ (a)); 3.33-3.22 (m; 2H; CH_2_(f)); 2.33 (s; 3H; CH_3_ (6)); 1.74-1.59 (m; 2H; CH_2_ (b)); 1.57-1.48 (m; 2H; CH_2_ (e)); 1.40-1.18 (m; 4H; CH_2_ (d/c)).

***(E)-N-(6-(5-(benzyloxy)-2-methyl-4-oxopyridin-1(4H)-yl)hexyl)-3-(3.4.5-trifluorophenyl)acrylamide (4c)***. Procedure C1. η = 48 %. ^1^H RMN (400 MHz, CDCl_3_): δ = 7.50-7.44 (m; 2H; CH (2’/6’)); 7.41 (d; *J* = 15.9 Hz; 1H; CH (β)); 7.37-7.27 (m; 5H; CH (3’-5’/2”/6”)); 6.57 (d; *J* = 15.7 Hz; CH (α)); 6.36 (s; CH (2)); 5.10 (s; CH (5)); 5.09 (s; 2H; CH_2_ (7)); 3.97-3.90 (m; 2H; CH_2_ (a)); 3.32-3.25 (m; 2H; CH_2_ (f)); 2.15 (s; 3H; CH_3_ (6)); 1.71-1.61 (m; 2H; CH_2_ (b)); 1.58-1.49 (m; 2H; CH_2_ (e)); 1.40-1.26 (m; 4H; CH_2_ (d/c)). ^13^C RMN (101 MHz, CDCl_3_): δ = 172.4 (C (4)); 166.2 (C (g)); 151.2 (ddd; *J*_CF_ = 248.8 Hz; *J* = 10.4 Hz; *J* = 4.2 Hz; C (3’’-5’’); 147.2 (C (3)); 146.8 (C (6)); <u>136.9</u> (C (β)); 136.5 (C (1’)); 131.9 (d; *J* = 5.3 Hz; C (1’’)) <u>128.1</u> (C (3’/5’)); <u>127.8</u> (C (2’/6’)); <u>127.7</u> (C (4’)); <u>127.5</u> ((2)); <u>123.4</u> (C (α)); <u>116.4</u> (C (5)); <u>111.5</u> (dd; *J*_CF_ = 16.3 Hz; *J*_CF_ = 5.6 Hz; C (2’’/6’’)); <u>70.9</u> (C (7)); <u>53.3</u> (C (a)); 39.0 (C (<u>f</u>)); <u>29.9</u> (C (e)); <u>28.8</u> (C (b)); <u>26.1</u> (C (c)); <u>25.5</u> (C (d)); <u>17.5</u> (C (8)).

***(E)-N-(6-(3-(benzyloxy)-2-methyl-4-oxopyridin-1(4H)-yl)hexyl)-3-(4-fluorophenyl)acrylamide (4d)***. Procedure C2. η = 50 %. ^1^H RMN (400 MHz, CDCl_3_): δ = 7.54 (d; *J* = 15.7 Hz; 1H; CH (β)); 7.48 (t; *J* = 5.7 Hz; 1H; NH); 7.45-7.40 (m; 2H; CH (2’’/6’’)); 7.38-7.34 (m; 2H; CH (2’/6’)); 7.32-7.26 (m; 3H; CH (3’-5’)); 7.20 (d; *J* = 7.5 Hz; 1H; CH (6)). 6.99-6.92 (m; 2H; CH (3’’/5’’)); 6.56 (d; *J* = 15.7 Hz; CH (α)); 6.37 (d; *J* = 7.5 Hz; 1H; CH (5)); 5.17 (s; 2H; CH (7)); 3.70 (t; *J* = 7.5 Hz; 2H; CH_2_ (a)); 3.34 (q; *J* = 6.5 Hz; 2H; CH_2_ (f)); 2.07 (s; 3H; CH_3_ (2)); 1.72-1.48 (m; 4H; CH_2_ (b/e)); 1.40-1.31 (m; 2H; CH_2_ (c)); 1.31-1.21 (m; 2H; CH_2_ (d)). ^13^C RMN (101 MHz, CDCl_3_): δ = 173.2 (C (4)); 166.3 (C (g)); 163.3 (d; *J*_CF_ = 249.7 Hz C (4’’); 146.1 (C (3)); 141.2 (C (2)); <u>138.7</u> (C (β)) <u>138.5</u> (C (6)); 137.3 (C (1’)); 131.4 (d; *J* = 3.2 Hz; (C (1’’)); <u>129.5</u> (d; *J* = 8.4 Hz; (C (2’’/6’’)); <u>128.9</u> (C (2’/6’)); <u>128.3</u> (C (3’/5’)); <u>128.1</u> (C (4’)); <u>121.5</u> (C (α)); <u>116.9</u> (C (5)); <u>115.7</u> (d; *J* = 21.7 Hz; (C (3’’/5’’)); <u>73.0</u> (C (7)); <u>53.8</u> (C (a)); 39.3 (C (f)); <u>30.5</u> (C (e)); <u>29.3</u> (C (b)); <u>26.4</u> (C (c)); <u>25.9</u> (C (d)); 12.4 (C (8)).

***(E)-N-(6-(3-(benzyloxy)-2-methyl-4-oxopyridin-1(4H)-l)hexyl)-3-(3.4-difluorophenyl)acrylamide (4e).*** Procedure C1. η = 63 %. ^1^H RMN (400 MHz, CDCl_3_): δ = 8.00 (t; *J* = 5.6; 1H; NH); 7.46 (d; *J* = 15.6 Hz; 1H; CH (β)); 7.37-7.32 (m; 2H; CH (2’/6’)); 7.30-7.22 (m; 5H; CH (3’-5’/6/6”)); 7.21-7.12 (m; 1H; CH (2”)); 7.05 (ddd; *J* = 10.0 Hz; *J* = 8.2 Hz; 1H; CH (5’’)); 6.62 (d; *J* = 15.6 Hz; CH (α)); 6.35 (d; *J* = 7.5 Hz; CH (5)); 5.16 (s; 2H; CH_2_ (7)); 3.71 (t; *J* = 7.5; 2H; CH_2_ (a)); 3.34 (q; *J* = 6.5 Hz; 2H; CH_2_ (f)); 2.08 (s; 3H; CH_3_ (2)); 1.67-1.50 (m; 4H; CH_2_ (b/e)); 1.41-1.31 (m; 2H; CH_2_ (c)); 1.31-1.23 (m; 2H; CH_2_ (d)). ^13^C RMN (101 MHz, CDCl_3_): δ = 173.1 (C (4)); 165.6 (C(g)); 150.9 (dd; *J*_CF_ = 248.6 Hz; *J*_CF_ = 13.2 Hz C (3’’); 150.5 (dd; *J*_CF_ = 248.6 Hz; *J*_CF_ = 13.2 Hz C (4”); 146.1 (C (3)); 141.1 (C (2)); <u>138.0</u> (C (β)); <u>138.2</u> (C (6)); 137.2 (C (1’)); 132.4 (dd; *J*_CF_ = 5.0 Hz; *J*_CF_ = 4.0 Hz C (1’’)); <u>129.1</u> (C (3’/5’)); <u>128.3</u> (C (2’/6’)); <u>128.1</u> (C (4’)); <u>124.6</u> (dd; *J*_CF_ = 6.3 Hz; *J*_CF_ = 3.3 Hz C (6’’)); <u>122.3</u> (C (α)); <u>117.6</u> (d; *J*_CF_ = 17.7 Hz; (C (5’’)); <u>117.1</u> (C (5)); <u>115.7</u> (d; *J*_CF_ = 17.6 Hz; (C (2’’)); <u>73.1</u> (C (7)); <u>53.9</u> (C (a)); 39.4 (C (<u>f</u>)); <u>30.5</u> (C (e)); <u>29.4</u> (C (b)); <u>26.4</u> (C (c)); <u>25.9</u> (C (d)); <u>12.4</u> (C (8)).

***(E)-N-(6-(3-(benzyloxy)-2-methyl-4-oxopyridin-1(4H)-yl)hexyl)-3-(3.4.5-trifluorophenyl)acrylamide (4f).*** Procedure C1. η = 32 %. ^1^H RMN (400 MHz, CDCl_3_): δ = 7.43 (d; *J* = 15.6 Hz; 1H; CH (β)); 7.39-7.36 (m; 2H; CH (2’/6’)); 7.39-7.36 (m; 4H; CH (3’-5’/6)); 7.15-7.03 (m; 2H; CH (2’’/6’’)); 6.79 (bs; 1H; NH); 6.48; (d; J = 15.6 Hz; CH (α)); 6.43 (d; *J* = 7.4 Hz; CH (5)); 5.20 (s; 2H; CH_2_ (7)); 3.78 (t; *J* = 7.5 Hz; 2H; CH_2_ (a)); 3.36 (q; *J* = 6.8 Hz; 2H; CH_2_ (f)); 2.11 (s; 3H; CH_3_ (2)); 1.68-1.60 (m; 2H; CH_2_ (b)); 1.61-1.51 (m; 2H; CH_2_ (e)); 1.43-1.24 (m; 4H; CH_2_ (d/c)). ^13^C RMN (101 MHz, CDCl_3_): δ = 172.9 (C (4)); 165.2 (C(g)); 151.3 (ddd; *J*_CF_ = 249.9 Hz; *J =* 10.1; *J =* 3.8 Hz C(3’’-5’’); 146.1 (C (3)); 141.3 (C (2)); <u>138.5</u> (C (6)); 137.3 (C (1’)); <u>137.1</u> (C (β)); 131.5 (d; *J =* 4.8; (C (1’’)); <u>129.0</u> (C (3’/5’)); <u>128.3</u> (C (2’/6’)); <u>128.1</u> (C (4’)); <u>123.83</u> (C (α)); <u>116.9</u> (C (5)); <u>111.7</u> (d; *J =* 5.8; (C(2’’)); <u>111.5</u> (d; *J =* 5.8; (C (6’’)); <u>73.1</u> (C (7)); <u>53.9</u> (C (a)); 39.4 (C (f)); <u>30.5</u> (C (e)); <u>29.23</u> (C (b)); <u>26.3</u> (C (c)); <u>25.9</u> (C (d)); <u>12.4</u> (C (8)).

##### 4.1.2.4. General procedures for the synthesis of final compounds **5a-f**

All final compounds **5a-f** were obtained following procedure **D**.

**Procedure D**. Compounds **4a-f** (1 mmol) were dissolved in a 1:1 TFA: Toluene solution (11 mL) and left to stir at 65 °C. After 24 hours, the solvents were evaporated under vacuum, and the resulting compound was washed in DCM and n-hexane to obtain a brownish oil. The experimental procedure was adapted from the literature [26].

(***E)-3-(4-fluorophenyl)-N-(6-(5-hydroxy-2-methyl-4-oxopyridin-1(4H)-yl)hexyl)acrylamide (5a)***. Procedure D. η = 85 %. ^1^H RMN (400 MHz, CDCl_3_): δ = 8.07 (s; 1H; CH (2)); 7.61-7.55 (m; 2H; CH (2’’/6’’))); 7.49 (d; *J* = 15.7 Hz; 1H; CH (β)); 7.10 (dd; *J =* 8.3 Hz; 2H; CH (3’’/5’’)); 7.06 (s; 1H; CH (5)); 6.57 (d; *J* = 15.7 Hz; CH (α)); 4.25 (d; *J* = 7.0 Hz. 2H; CH_2_ (a)); 3.37-3.25 (m. 2H; CH_2_ (f)); 2.59 (s; 3H; CH_3_ (6)); 1.85 (bs. 2H; CH_2_ (b)); 1.60 (bs. 2H; CH_2_ (e)); 1.42 (bs; 4H; CH_2_ (d/c)). ^13^C RMN (101 MHz, CDCl_3_): δ = 167.1 (C (4)); 163.5 (d; *J*_CF_ = 248.7 Hz; C (4’’); 160.9 (C(g)); 147.4 (C (3)); 144.5 (C (6)); <u>138.9</u> (C (β)); 131.4 (d; *J* = 3.3 Hz; C (1’’)); <u>130.4</u> (C (2)); <u>129.5</u> (d; *J* = 8.4 Hz; C (2’’/6”)); <u>120.6</u> (C (α)); <u>115.6</u> (d; *J* = 22.1 Hz; C (3’’/5’’)); <u>114.0</u> (C (5)); <u>55.6</u> (C (a)); <u>38.9</u> (C (f)); <u>29.8</u> (C (e)); <u>28.8</u> (C (b)); <u>26.0</u> (C (c)); <u>25.5</u> (C (d)); <u>17.7</u> (C (8)).

***(E)-3-(3.4-difluorophenyl)-N-(6-(5-hydroxy-2-methyl-4-oxopyridin-1(4H)-yl)hexyl)acrylamide (5b).*** Procedure D. η = 56 %. ^1^H RMN (400 MHz, CDCl_3_): δ = 8.02 (s; 1H; CH (2)); 7.55-7.45 (m; 1H; CH (6’’))); 7.45 (d; *J* = 16.0 Hz; 1H; CH (β)); 7.39-7.34 (m; 1H; CH (5’’)); 7.32-7.22 (m; 1H; CH (2’’)); 6.98 (s. 1H; CH (5); 6.56 (d; *J* = 15.7 Hz; CH (α)); 4.53-4.01 (m; 2H; CH_2_ (a)); 3.36-3.26 (m; 2H; CH_2_ (f)); 2.57 (s; 3H; CH_3_ (6)); 1.85 (bs; 2H; CH_2_ (b)); 1.66-1.54 (m; 2H; CH_2_ (e)); 1.50-1.43 (m; 4H; CH_2_ (d/c)). ^13^C RMN (101 MHz, CDCl_3_): δ = 166.7 (C (4)); 161.9 (C (g)); 151.0 (dd; *J*_CF_ = 250.3 Hz; *J*_CF_ = 12.9 Hz; C (3’’); 150.4 (dd; *J*_CF_ = 247.2 Hz; *J*_CF_ = 13.0 Hz; C (4’’); 147.3 (C (3)); 144.8 (C (6)); <u>137.9</u> (C (β)); 132.6 (dd; *J*_CF_ = 6.1 Hz; *J*_CF_ = 3.9 Hz; C (1’’)); <u>129.8</u> (C (2)); <u>124.5</u> (dd; *J*_CF_ *=* 6.6 Hz; *J*_CF_ = 3.4 Hz; C (6’’)); <u>121.9</u> (C (α)); <u>117.4</u> (d; *J* = 17.9 Hz; C (5’’)); <u>115.6</u> (d; *J* = 17.9 Hz; C(2’’)); <u>114.0</u> (C (5)); <u>55.3</u> (C (a)); <u>38.9</u> (C (f)); <u>29.8</u> (C (e)); <u>28.8</u> (C (b)); <u>26.0</u> (C (c)); <u>25.5</u> (C (d)); <u>17.7</u> (C (8)).

***(E)-3-(3.4.5-trifluorophenyl)-N-(6-(5-hydroxy-2-methyl-4-oxopyridin-1(4H)-yl)hexyl)acrylamide (5c)***. Procedure D. η = 86 %. ^1^H RMN (400 MHz, CDCl_3_): δ = 8.02 (s; 1H; CH (2)); 7.41 (d; *J* = 16.0 Hz; 1H; CH (β)); 7.36 (dd; *J* = 8.8 Hz; *J* = 6.7 Hz 2H; CH (2’’/6”))); 6.99 (s; 1H; CH (5)); 6.57 (d; *J* = 15.7 Hz; CH (α)); 4.27-4.20 (m; 2H; CH_2_); 3.33-3.28 (m; 2H; CH_2_ (f)); 2.58 (s; 3H; CH_3_ (6)); 1.85 (bs; 2H; CH_2_ (b)); 1.64-1.54 (m. 2H; CH_2_ (e)); 1.55-1.39 (m; 4H; CH_2_ (d/c)). ^13^C RMN (101 MHz, CDCl3): δ = 166.2 (C (4)); 162.1 (C (g)); 151.3 (ddd; *J*_CF_ = 248.5 Hz; *J* = 10.1 Hz; *J* = 3.7 Hz; C (3’’-5’’); 147.3 (C (3)); 144.8 (C (6)); <u>136.9</u> (C (β)); 131.9 (d; *J* = 4.8 Hz; C (1’’)); <u>129.7</u> (C (2)); <u>123.4</u> (C (α)); <u>114.0</u> (C (5)); <u>111.5</u> (dd; *J* = 16.6 Hz*; J* = 6.1 Hz; C(2’’/6”)); <u>55.3</u> (C (a)); <u>38.9</u> (C (f)); <u>29.8</u> (C (e)); <u>28.8</u> (C (b)); <u>26.0</u> (C (c)); <u>25.0</u> (C (d)); <u>17.6</u> (C (8)).

***(E)-3-(4-fluorophenyl)-N-(6-(3-hydroxy-2-methyl-4-oxopyridin-1(4H)-yl)hexyl)acrylamide (5d).*** Procedure D. η = 47 %. ^1^H RMN (400 MHz, CDCl_3_): δ = 7.92 (d; *J* = 6.7 Hz; 1H; CH (6)); 7.58 (dd; *J* = 8.4 Hz; *J* = 5.5 Hz; 2H; CH (2’’/6’’))); 7.50 (d; *J* = 15.6 Hz; 1H; CH (β)); 7.12 (dd; *J* = 8.6 Hz; 2H; CH (3’’/5’’)); 6.81 (d; *J* = 6.7Hz; 1H; CH (5)); 6.53 (d; *J* = 15.7 Hz; CH (α)); 4.23 (t; *J* = 7.6 Hz. 2H; CH_2_ (a)); 3.33-3.28 (m; 2H; CH_2_ (f)); 2.55 (s; 3H; CH_3_ (2)); 1.90-1.73 (m; 2H; CH_2_ (b)); 1.66-1.53 (m;; 2H; CH_2_); 1.49-1.38 (m; 4H; CH_2_ (d/c)). ^13^C RMN (101 MHz, CDCl_3_): δ = 167.1 (C (4)); 164.8 (C(g)); 162.6 (ddd; *J*_CF_ = 39.5 Hz; C(4’’); 144.6 (C (3)); <u>138.91</u> (C (β)); <u>137.5</u> (C (6)); 131.3 (d; *J* = 3.5 Hz; (C (1’’)); <u>129.5</u> (d; *J* = 8.5 Hz; (C (2’’/6’’)); <u>120.4</u> (C (α)); <u>115.4</u> (d; *J* = 22.2 Hz; (C (3’’/5’’)); <u>110.7</u> (C (5)); <u>55.32</u> (C (a)); 38.9 (C (<u>f</u>)); <u>30.0</u> (C (e)); <u>28.9</u> (C); <u>26.0</u> (C (c)); <u>25.5</u> (C (d)); <u>10.9</u> (C (8)).

***(E)-3-(3.4-difluorophenyl)-N-(6-(3-hydroxy-2-methyl-4-oxopyridin-1(4H)-yl)hexyl)acrylamide (5e)***. Procedure D. η = 68 %. ^1^H RMN (400 MHz, CDCl_3_): δ = 8.13 (d; *J* = 7.0 Hz; 1H; CH (6)); 7.54-7.47 (m; 2H; CH (6’’))); 7.45 (d; *J* = 16.0 Hz; 1H; CH (β)); 7.40-7.33 (m; 1H; CH (5’’)); 7.32-7.23 (m; 1H; CH (2’’)); 7.10 (d; J = 6.9 Hz; CH(5)); 6.56 (d; *J* = 15.7 Hz; CH (α)); 4.40-4.30 (m; 2H; CH_2_ (a)); 3.33-3.28 (m; 2H; CH_2_ (f)); 2.63 (s; 3H; CH_3_ (2)); 1.85 (m; 2H; CH_2_ (b)); 1.60 (m; 2H; CH_2_ (e)); 1.45 (m; 4H; CH_2_ (d/c)). ^13^C RMN (101 MHz, CDCl_3_): δ = 166.7 (C (4)); 158.4 (C(g)); 150.9 (dd; *J*_CF_ = 250.4 Hz; *J*_CF_ *=* 12.7 Hz; C(3’’); 150.4 (dd; *J*_CF_ = 247.2 Hz; *J*_CF_ *=* 13.1 Hz; C(4”); 143.8 (C (3)); 141.5 (C (2)); <u>137.9</u> (C (β)); <u>137.6</u> (C (6)); 132.6 (dd; *J*_CF_ = 6.2 Hz; *J*_CF_ *=* 4.0 Hz (C (1’’)); <u>124.5</u> (dd; *J*_CF_ = 6.5 Hz; *J*_CF_ *=* 3.4 Hz (C (6’’)); 121.9 (C (α)); <u>117.4</u> (d; *J =* 17.8 Hz (C (5’’)); <u>115.6</u> (d; *J =* 17.8 Hz (C(2’’)); <u>110.3</u> (C (5)); <u>56.4</u> (C (a)); 38.9 (C (e)); <u>29.8</u> (C (e)); <u>28.8</u> (C (b)); <u>25.9</u> (C (c)); <u>25.5</u> (C (d)); <u>11.3</u> (C (8)).

***(E)-3-(3.4.5-trifluorophenyl)-N-(6-(3-hydroxy-2-methyl-4-oxopyridin-1(4H)-yl)hexyl)acrylamide (5f).*** Procedure D. η = 86 %. ^1^H RMN (400 MHz, CDCl_3_): δ = 8.14 (d; *J* = 6.9 Hz; 1H; CH (6)); 7.39 (d; *J* = 16.5 Hz; 1H; CH (β)); 7.37-7.32 (m; 2H; CH (2”/6’’))); 7.15 (d; *J* = 6.9 Hz; 1H; CH(5)); 6.61 (d; *J* = 15.7 Hz; CH (α)); 4.37 (t; *J* = 7.7 Hz; 2H; CH_2_ (a)); 3.35-3.28 (m; 2H; CH_2_ (f)); 2.64 (s; 3H; CH_3_ (2)); 1.93-1.84 (m; 2H; CH_2_ (b)); 1.67-1.56 (m; 2H; CH_2_ (e)); 1.50-1.39 (m; 4H; CH_2_ (d/c)). ^13^C RMN (101 MHz, CDCl_3_): δ = 166.3 (C (4)); 158.5 (C(g)); 151.2 (ddd; *J*_CF_ = 248.8 Hz; *J*_CF_ = 10.1 Hz; *J*_CF_ = 3.7 Hz; C(3’’-5’’); 143.8 (C (3)); 141.3 (C (2)); 139.7 (ddd; *J*_CF_ = 253.5 Hz; *J*_CF_ = 16.1 Hz; (C(4”)); <u>137.6</u> (C (6)); <u>136.9</u> (C (β)); 132.0-131.7 (m; (C (1’’)); <u>123.4</u> (C (α)); <u>111.5</u> (d; *J*_CF_ = 16.0; *J*_CF_ = 5.8 Hz; (C(2’’)); <u>111.6</u> (d; *J*_CF_ = 16.0; *J*_CF_ = 5.8 Hz; (C (6’’)); <u>110.4</u> (C (5)); <u>56.4</u> (C (a)); 39.0 (C (<u>f</u>)); <u>29.8</u> (C (e)); <u>28.7</u> (C (b)); <u>25.9</u> (C (c)); <u>25.5</u> (C (d)); <u>11.4</u> (C (8)).

### 4.2 Antimicrobial screening

Antimicrobial screening was conducted by the Community for Open Antimicrobial Drug Discovery (CO-ADD, Australia). The compounds were evaluated against a panel of clinically relevant microorganisms, including methicillin-resistant *S. aureus* ATCC 43300 (MRSA), E. coli ATCC 25922, *K. pneumoniae* ATCC 700603, *P. aeruginosa* ATCC 27853, and *A. baumannii* ATCC 19606, along with two yeast strains *C. albicans* ATCC 90028 and *Cryptococcus neoformans* ATCC 208821. The bacterial strains were cultured in cation-adjusted mueller hinton broth (CAMHB; Bacto Laboratories 212322) at 37 °C overnight. Before testing, a sample of each culture was diluted 40-fold in fresh broth and incubated at 37°C for 1.5-3 h to reach the mid-log phase. The resultant cultures were diluted with CAMHB (as determined by OD_600_) and plated, giving a final cell density of 5 x 10^6^ CFU/mL and a total volume of 50 µL. Plates were incubated at 37 °C for 18 h without shaking and the growth inhibition was determined by measuring the absorbance at 600 nm using a Tecan M1000 Pro monochromator plate reader.

The fungi strains were cultured on yeast extract-peptone dextrose (YPD; Becton Dickinson 242720) agar at 30°C for three days. After this period, a yeast suspension of 2.5 x 10^3^ CFU/mL (as determined by OD) was prepared from five colonies on yeast nitrogen base (YNB, Becton Dickinson 233520) and a total volume of 50 µL was added to each well containing the compounds. Plates were incubated at 35 °C for 24 h without shaking. Growth inhibition of *C. albicans* was quantified by measuring the absorbance at 530 nm (OD_530_). Whereas for *C. neoformans*, resazurin was first added (0.001% final concentration) and the plates incubated at 35 °C for 2 h. Then, the growth inhibition was determined by measuring the difference in absorbance between 600 and 750 nm (OD_600-570_), using a Biotek Synergy HTX plate reader.

Colistin sulfate (Sigma Aldrich, Castle Hill, Australia C4461) and vancomycin hydrochloride (Sigma Aldrich, 861987) were used as positive bacterial inhibitor controls for Gram-negative and Gram-positive bacteria, respectively. While for *C. albicans* and *C. neoformans*, the positive fungal inhibitor control used was fluconazole (Sigma Aldrich, F8929). Stock solutions from the compounds under study were prepared in DMSO to a final concentration of 32 µg/mL. The percentage of DMSO was kept to a maximum of 1% DMSO and the solutions were stored frozen at –20 °C. In the whole cell inhibition assays, compounds were prepared in 384-well non-binding surface (NBS) plates for each bacterial/fungal strain as a 2-fold dose response from 32 to 0.25 µg/mL, with a maximum concentration of DMSO of 0.5%. The sample preparation was done using liquid handling robots.

### 4.3. Haemolysis assays

Haemolytic activity was evaluated using human red blood cells (sourced from LifeBlood) in collaboration with CO-ADD (Australia). The use of human blood was approved by the University of Queensland Institutional Human Research Ethics Committee, Approval Number 2020001239. Briefly, human whole blood was washed three times with three volumes of 0.9% NaCl and resuspended to a concentration of 0.5 × 10^8^ cells/mL. The washed cells were counted manually in a Neubauer haemocytometer and plated in the polypropylene 384-well plates (Corning, 3657) containing the compounds for a final volume of 50 µL. The plates were left shaking for 10 min and then incubated at 37 °C for 1 h. After this period, the plates were centrifuged for 10 min at 1000 × g to pellet cells and debris and then 25 µL of the supernatant was transferred to reading plates. Haemolysis was evaluated by measuring the supernatant absorbance at 405 nm (OD_405_), using a M1000 Pro monochromator plate reader (Tecan Trading AG, Switzerland). HC_10_ values were obtained from dose-response curves, with variable fitting values for bottom, top and slope. The curve fitting was determined using Pipeline Pilot’s dose–response component to calculate the HC_50_ values, which were then converted into HC_10_ values using **Equation (1)**.

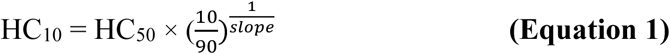

Melittin was used as positive reference. Stock solutions were prepared in DMSO to a final concentration of 1% (v/v) by liquid handling robots, with a maximum concentration of 1% DMSO (v/v) and stored frozen at –20 °C.

### 4.4. Cytotoxicity

#### 4.4.1. Reagents and materials

The neutral red solution, resazurin and trypan blue solution 0.4 % (w/v) were purchased from Sigma. Fetal bovine serum (FBS) and Dulbecco’s Modified Eagle Medium (DMEM) with a high glucose content were obtained from Gibco Laboratories. Phosphate buffered saline (PBS) was purchased from Alfa Aesar. Hank’s balanced salt solution (HBSS) without Ca^2+^ and Mg^2+^ and the antibiotic (penicillin 10.000 U/mL, streptomycin 10.000 µg/mL) were obtained from Pan Biotech. Trypsin (0.25 %) was purchased from Speciality media. Dimethyl sulphoxide (DMSO), methanol, 70 % ethanol and acetic acid were obtained from Merck.

The cell culture was carried out in a Haier HR 1200-IIA2-D Biological Safety Cabinet. All the solutions used for cell culture were heated to 37 °C in a Thermo Scientific Precision GP 20 water bath. The cells used in this work were maintained in a Thermo Scientific HERAcell 150i incubator at 37 °C with 5 % CO_2_. The cells were counted using an Invitrogen Countess 3 high-performance automated cell counter. The absorbance and fluorescence readings were taken on a Biotek Synergy HT microplate reader and 96-well plates from Techno Plastic Products.

#### 4.4.2. Cytotoxicity screening assay using HEK293 cells

The initial cytotoxicity screening was performed by CO-ADD using human embryonic kidney cells (HEK293), a cell line widely employed in preclinical studies to assess the drug’s safety profile. HEK293 cells (ATCC CRL-1573) were plated in 384-well tissue-culture treated black plates (TC-treated; Corning, 3712/3764) containing the compounds to give a density of 5000 cells/well in a final volume of 50 µL, as determined by manual cell count in a Neubauer haemocytometer. DMEM supplemented with 10% FBS was used as growth media and the cells were incubated with the compounds for 20 h at 37 °C in 5% CO_2_. After the addition of resazurin (25 µg/mL final concentration) and incubation at 37 °C and 5% CO_2_ for additional 3 h, the growth inhibition of HEK293 cells was determined by recording the fluorescence at ex:530/10 nm and em:590/10 nm (F560/590). The fluorescence was measured using a Tecan M1000 Pro monochromator plate reader. The percentage of growth inhibition was calculated for each well, using the negative control (media only) and positive control (cell culture without inhibitors) on the same plate as references. CC_50_ values were obtained from dose-response curves, with variable fitting values for bottom, top and slope. The curve fitting was implemented using Pipeline Pilot’s dose–response component.

Tamoxifen was used as positive cytotoxic reference. Stock solutions were prepared in DMSO to a final concentration of 1% (v/v) by liquid handling robots, with a maximum concentration of 1% DMSO (v/v) and stored frozen at –20 °C.

#### 4.4.3. Cell viability assays using HepG2 cells

The cytotoxicity of the final **5a-f** compounds was assessed using a cell line with high mitochondrial expression, namely human hepatocellular carcinoma cells (HepG2). The cells were maintained in culture with DMEM medium supplemented with 10 % FBS (v/v), penicillin 10.000 U/mL and streptomycin 10 mg/mL at 37 °C with 5 % CO_2_. The cell line was routinely harvested into new 75 cm^2^ culture flasks by trypsinization when confluence reached 70-80 %. For trypsinization, the cells were washed with PBS and treated with trypsin (0.25 %) for 5 min at 37 °C. After this, the cells were resuspended in the respective cell medium, and the live cells were counted using trypan blue. After counting, the cells were cultured in 96-well plates at a density of 6.25 × 10^4^ cells/cm^2^, equivalent to 2.0 × 10^5^ cells/mL. Exposure to the compounds was carried out 24 hours after cell plating with the same cell medium.

The experimental procedure was adapted from the literature [27, 28].

##### 4.4.3.1. Neutral red assessment

In this assay, HepG2 cells were treated after adhesion with DMEM medium in the presence or absence of the **5a-f** compounds at different concentrations (0.1, 0.5, 1. 5, 10 and 25 µM) in transparent 96-well plates and incubated at 37 °C with 5 % CO_2_ for 24 h. The control, which consisted of medium and DMSO (0.1 %) was also placed in 6 wells. Briefly, after the incubation time, the cells were incubated with the NR solution for 1 hour to 1 hour and 30 minutes. This solution consists of a dilution of a stock solution of neutral red (3.3 mg/mL in medium). After this time, the cells were incubated with a lysis solution at room temperature and shaken until all the wells were uniform. This lysis solution consists of a mixture of absolute ethanol/distilled water (1:1) containing 5% glacial acetic acid to extract the dye that has accumulated in the lysosomes of viable cells. The NR extracted from the cells was measured on a microplate reader at 540 nm.

##### 4.4.3.2. Resazurin assessment

In this test, HepG2 cells were treated after adhesion with DMEM medium in the presence or absence of the **5a-f** compounds at different concentrations (0.1, 0.5, 1. 5, 10 and 25 µM) in transparent 96-well plates and incubated at 37 °C with 5 % CO_2_ for 24 h. Controls were also placed in 6 wells. Briefly, after the incubation time, the cells were subjected to resazurin solution for 1 hour. This solution consists of a dilution of a stock solution of resazurin (1.1 mg/mL in medium). After this time, the resazurin extracted from the cells was measured on a microplate reader at 540 nm.

##### 4.4.3.3. Statistical Analysis and Software

Data were evaluated using the one-way analysis of variance (ANOVA) test followed by Dunnett’s test for multiple comparisons. The results were expressed as mean and standard deviation (mean ± DP (standard deviation) of a minimum of three independent experiments performed in triplicate. Values of p < 0.05 were considered statistically significant. All statistical analysis was performed using GraphPad Prism Software 8.0.

### 4.5. Determination of the chromatographic hydrophobicity index

Lipophilicity was determined from CHI (Chromatographic Hydrophobicity Index) values, which are based on the retention time of compounds when passing through a reversed stationary phase using HPLC and can be calculated using the following equation:

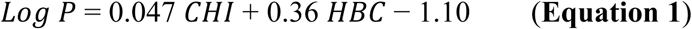

where HBC corresponds to the number of hydrogen-bond donors.

CHI values were obtained from the experimental retention times (tr) of the target compounds and a mixture of reference standards, following a previously reported protocol at pH 2.6 [29]. Analyses were performed using an HPLC system equipped with a Luna C18(2) column (Phenomenex, CA, USA; 150 × 4.6 mm, 5 µm). Compounds **5a–f** were prepared from 50 mM stock solutions in DMSO and diluted to a final concentration of 125 µM in acetonitrile:water (1:1, v/v). Mobile phase A was aqueous acetic acid 1% (v/v) (pH 2.6), and mobile phase B was acetonitrile. The following gradient conditions were also applied: flow rate of 1 mL/min, injection volume of 20 µL and solvent B gradient of 0-1 min 0%, 1-7 min 0 –>100%, 7-10 min 100%, 10-12 min 100% –> 0%, and 13 min 0%. A calibration curve was obtained using a mixture of reference compounds (**Figure 5**). The results are expressed as mean values of three experiments (**Table 4)**.

**Figure 5.**
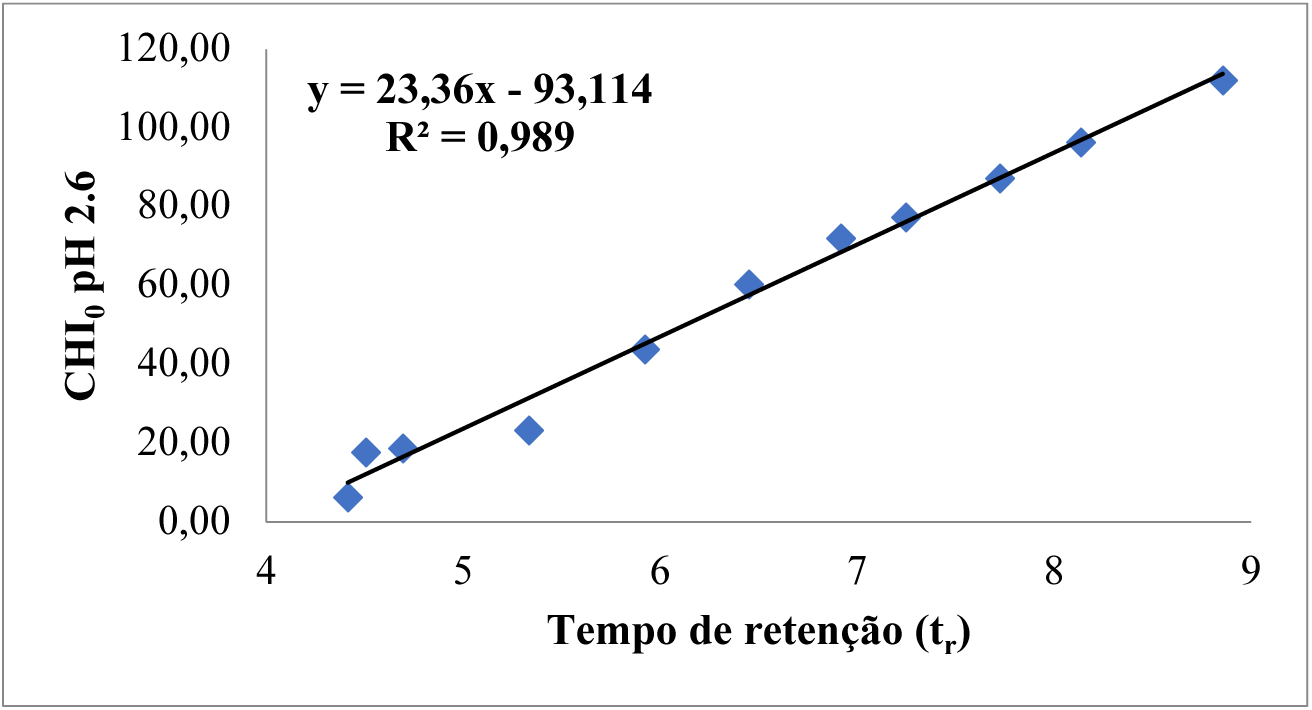
Calibration curve of the CHI value as a function of retention time at pH 2.6.

**Table 4.**
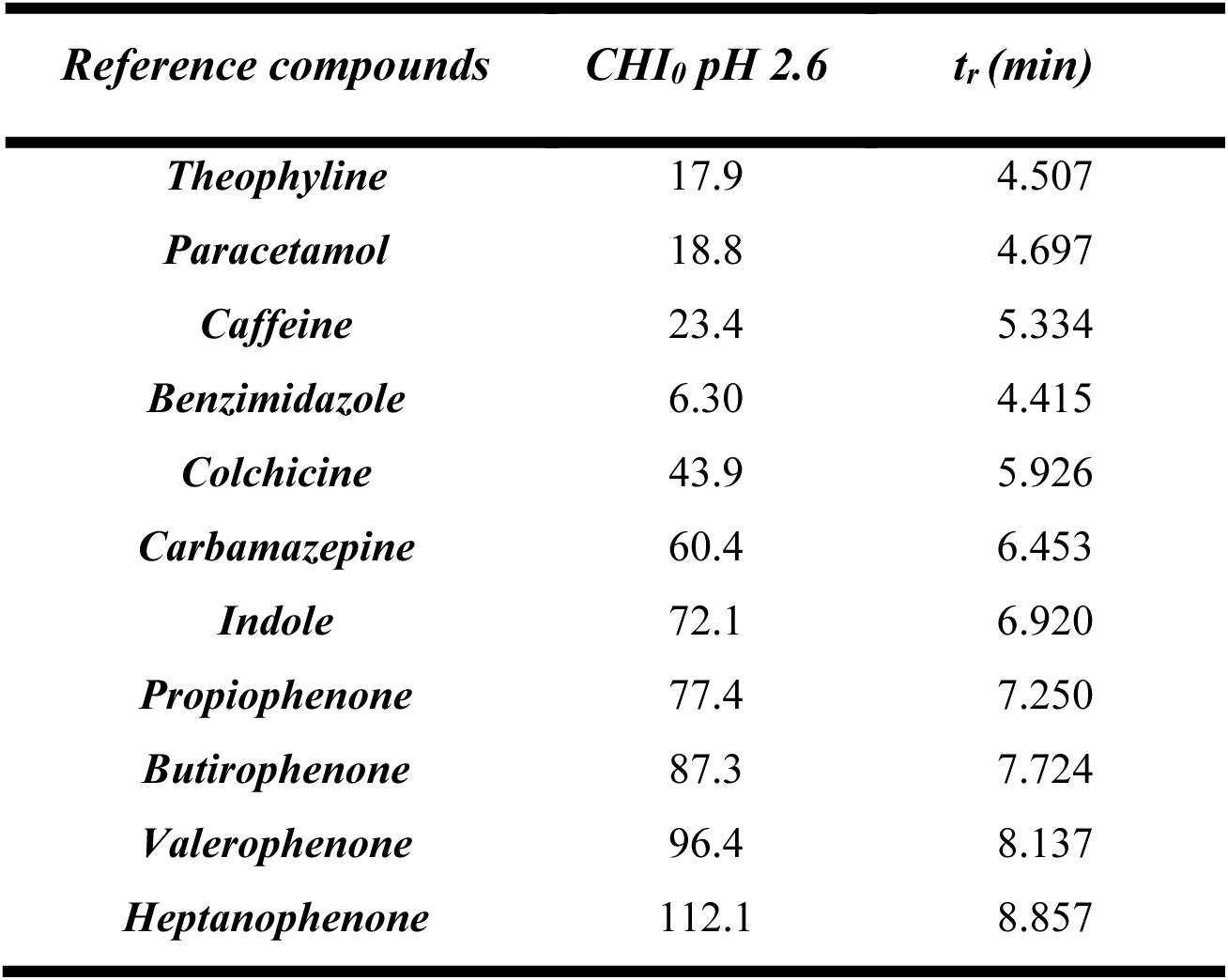
Retention time and respective CHI values of the reference compounds at pH 2.6.

### 4.6. Assessment of metal chelation properties

This study was carried out using UV-Vis spectrometry. The assay was performed on a microplate, and all determinations were carried out in triplicate. First. a solution of the compounds to be tested (40 µM) with or without Fe (III) (20 µM) in phosphate-buffered saline (PBS) (1×. pH 7.4) was added to each well of the microplate. with the plate been incubated for 30min at temperature environment.

Absorption spectra were obtained between 200-550 nm with 2 nm intervals. Next. the stoichiometry of the compound-Fe(III) complex was calculated following the molar ratio method. For this. a solution of each compound (40 µM in DMSO) was incubated with increasing concentrations of Fe(III) chloride (0-100 µM in EtOH) in PBS and the absorption spectrum was recorded. PBS. EtOH. and DMSO were used as blanks.

The procedure was adapted from the literature [30].

## Abbreviations

AMR, antimicrobial resistance; Boc, Tert-butyloxycarbonyl; CHI, Chromatographic Hydrophobicity Index; DCM, dichloromethane; DMEM, Dulbecco’s Modified Eagle Medium; DMSO, Dimethylsulfoxide; FBS, Fetal Bovine serum; HBSS, Hanks’ Balanced Salt Solution; HPLC, High-performance liquid chromatography; MeOD, Deuterated methanol; NR, Neutral Red; OMS, World Health Organization; p.a. Quality for analysis; PBS, Phosphate Buffered Saline; RMN, Nuclear Magnetic Resonance; TFA, Trifluoroacetic acid; TLC, Thin Layer Chromatography; TMS, Tetramethylsilane; tr, retention time; UV, ultraviolet.

## Acknowledgments

This work was funded by FEDER funds through the Operational Program Competitiveness Factors COMPETE and national funds by the FCT-Foundation for Science and Technology under research grant UID/BIM/04308/2019. This work was also funded by project IMPULSE: IMPROVING USER EXPERIENCE, LONG-TERM SUSTAINABILITY, AND SERVICES OF EU-OPENSCREEN (Funding programme: Horizon Europe; Grant agreement number: 101132028). S.B. (2023.06106.CEECIND) contract was supported by FCT and FEDER/COMPETE funds.

The authors thank CO-ADD (The Community for Open Antimicrobial Drug Discovery) not-for-profit initiative, funded by the Welcome Trust (UK) and The University of Queensland (Australia) for performing the preliminary antimicrobial and toxicity screenings.

